# Identification of chromatin states during zebrafish gastrulation using CUT&RUN and CUT&Tag

**DOI:** 10.1101/2021.06.22.447589

**Authors:** Bagdeser Akdogan-Ozdilek, Katherine L Duval, Fanju W Meng, Patrick J Murphy, Mary G Goll

**Author notes:** authors contributed equally (Bagdeser Akdogan-Ozdilek, Katherine L Duval). co-corresponding authors Contacts: Mary Goll, 120 E Green Street, Athens GA, 30602, Patrick Murphy, 601 Elmwood Ave, Rochester, NY, 14642.

## Abstract

**Background:** Cell fate decisions are governed by interactions between sequence-specific transcription factors and a dynamic chromatin landscape. Zebrafish offer a powerful system for probing the mechanisms that drive these cell fate choices, especially in the context of early embryogenesis. However, technical challenges associated with conventional methods for chromatin profiling have slowed progress toward understanding the exact relationships between chromatin changes, transcription factor binding, and cellular differentiation during zebrafish embryogenesis.

**Results:** To overcome these challenges, we adapted the chromatin profiling methods CUT&RUN and CUT&Tag for use in zebrafish, and applied these methods to generate high resolution enrichment maps for H3K4me3, H3K27me3, H3K9me3, RNA polymerase II, and the histone variant H2A.Z from mid gastrula stage embryos. Using this data, we identify a conserved subset of developmental genes that are enriched in both H3K4me3 and H3K27me3 during gastrulation, provide evidence for an evolving H2A.Z landscape during embryo development, and demonstrate the increased effectiveness of CUT&RUN for detecting protein enrichment at repetitive sequences.

**Conclusions:** Our results demonstrate the power of combining CUT&RUN and CUT&Tag methods with the strengths of the zebrafish system to define emerging chromatin landscapes in the context of vertebrate embryogenesis.

## Introduction

Interaction between proteins and DNA is central to regulation of transcription and generation of the complex gene expression patterns required for normal development. Transcription factor binding helps to control the rate of gene expression in a given tissue, while modification of the histones that package DNA creates localized chromatin environments that promote or repress transcription factor binding (Lawrence et al., 2016; Talbert et al., 2019). Modification of histone tails can be highly dynamic, driving dramatic reshaping of chromatin, especially during early animal embryogenesis. For example, in zebrafish, large scale *de novo* establishment of histone lysine 4 trimethylation (H3K4me3), histone lysine 27 trimethylation (H3K27me3) and histone lysine 9 trimethylation (H3K9me3) is first noted during blastula stages, with genome-wide accumulation of these modifications loosely coinciding with zygotic genome activation (Akdogan-Ozdilek et al., 2020; Horsfield, 2019). Similar dynamic changes in histone modifications have been noted during early embryogenesis in mouse, Drosophila, and *C.* elegans (Schulz and Harrison, 2019; Wu and Vastenhouw, 2020; Xu et al., 2021).

For roughly three decades, Chromatin Immunoprecipitation (ChIP) has been the primary method used to detect protein-DNA interactions (Solomon and Varshavsky, 1985). In its standard form, this method requires crosslinking of proteins to DNA, physical fragmentation of chromatin, and antibody mediated selection of chromatin fragments of interest. ChIP has proven especially powerful when coupled with high throughput sequencing (ChIP-seq) and has been widely implemented to profile genome-wide chromatin states and transcription factor binding (Barski et al., 2007; Johnson et al., 2007; Park, 2009). However, the need for crosslinking, relatively low signal to noise ratios, and input requirements of ~1 million cells present challenges to effective implementation of ChIP, especially when applied to early-stage embryos and embryonic tissues. Modifications of conventional protocols that allow ChIP to be performed in the absence of crosslinking and with low cell numbers have been described (Brind’Amour et al., 2015; Kasinathan et al., 2014). However, these methods can require deep sequencing, and generally work best for abundant proteins.

Cleavage Under Targets and Release Using Nuclease (CUT&RUN) offers a recently described alternative to ChIP that may be especially appealing for studies requiring early-stage embryos and embryonic tissues (Meers et al., 2019; Skene and Henikoff, 2017). Based on Chromatin Immunocleavage (ChIC) technology (Schmid et al., 2004), CUT&RUN employs a targeted nuclease strategy to isolate DNA fragments associated with proteins of interest. In CUT&RUN, a fusion of micrococcal nuclease and protein A/G (pAG-MNase) selectively cleaves antibody-bound chromatin in unfixed permeabilized cells. Released fragments are then isolated from solution and used for sequencing. The strategy bypasses immunoprecipitation steps, thereby shortening time required for purification. By only releasing the relevant fraction of the genome, CUT&RUN promotes high signal-to-noise ratios that allow significantly lower sequencing read depths (1/10th that of ChIP) and use of far fewer cells for analysis (1/10 to 1/1000th of ChIP) (Meers et al., 2019; Skene and Henikoff, 2017). Compared to physical fragmentation, endonuclease digestion also generates shorter fragments that reflect true protein footprints, allowing for base pair resolution mapping of protein binding in the absence of crosslinking.

Cleavage Under Targets and Tagmentation (CUT&Tag) is a recently developed variation of CUT&RUN that also takes advantage of immunotethering to identify regions of antibody-bound chromatin (Kaya-Okur et al., 2019). However, in this method, a fusion of protein A/G and Tn5 transposase is used to catalyze simultaneous cleavage and sequencing adapter ligation at antibody-bound chromatin, thus eliminating the need for downstream library preparation. Similar to CUT&RUN, this streamlined approach yields improved signal-to-noise ratios, especially when mapping histone modifications using ultra-low sample inputs (Kaya-Okur et al., 2019).

To date, CUT&RUN has been used primarily to profile transcription factor binding and histone modification enrichment in mammalian cells (Daneshvar et al., 2020; Federation et al., 2020; Hainer and Fazzio, 2019; Hyle et al., 2019; Kong et al., 2021; Liu et al., 2018; Meers et al., 2019; Oomen et al., 2019; Pagin et al., 2021; Park et al., 2019; Pritykin et al., 2021; Roth et al., 2018; Skene and Henikoff, 2017). Success has also been reported adapting CUT&RUN for use in S. cerevisiae (Skene and Henikoff, 2017), Drosophila (Ahmad and Spens, 2019; Uyehara and McKay, 2019), Arabidopsis (Zheng and Gehring, 2019), mouse germ cells (Ernst et al., 2019; Menon et al., 2019; Prakash and Barau, 2021) and mouse and human embryos (Inoue et al., 2018; Zhang et al., 2019). Thus far, CUT&Tag has been applied to assay enrichment of modified histones in plant and mammalian tissues (Bartosovic et al., 2021; Kaya-Okur et al., 2019; Murphy et al., 2020; Tao et al., 2020; Wang et al., 2021; Wu et al., 2021; Yu et al., 2021).

Here we describe successful implementation of CUT&RUN and CUT&Tag methods for profiling protein-DNA interactions in zebrafish embryos. Zebrafish have long been recognized as a powerful system for the study of early vertebrate development, as external fertilization facilitates molecular analysis at the earliest stages of embryogenesis. However, classical ChIP experiments in zebrafish can still require hundreds of embryos, resolution is limited, and developmentally emerging sites of enrichment can be difficult to identify above background. We apply modified CUT&RUN and CUT&Tag methods to generate high resolution maps of enrichment for H3K4me3, H3K27me3, H3K9me3, RNA polymerase II (pol II) and the histone variant H2A.Z during zebrafish gastrulation. Using this data, we identify a conserved subset of developmental genes that are enriched in H3K4me3 and H3K27me3 during gastrulation, provide evidence for a changing H2.AZ landscape during embryogenesis, and demonstrate increased effectiveness of CUT&RUN for detecting protein enrichment at repetitive sequences with reduced mappability. Our work demonstrates the power of combining CUT&RUN and CUT&Tag with the strengths of the zebrafish system to better understand the changing embryonic chromatin landscape and its roles in shaping development.

## Results

### CUT&RUN effectively detects H3K4me3 and H3K27me3 enrichment near gene transcriptional start sites (TSS)

To test the feasibility of using CUT&RUN in zebrafish, we initially profiled two well-characterized histone modifications, H3K4me3 and H3K27me3, in mid-gastrula stage embryos (shield stage, ~6 hours post fertilization (hpf)). H3K4me3 is typically associated with the TSS of active genes, while H3K27me3 associates with repressed genes. The combined presence of H3K4me3 and H3K27me3 is also observed at a subset of gene promoters during embryogenesis, with these bivalently marked genes existing in a poised state that enables rapid activation at appropriate developmental time points (Bernstein et al., 2006). Previous studies have identified gene regulatory regions marked by both H3K27me3 and H3K4me3 in blastula stage zebrafish embryos (Lindeman et al., 2011; Vastenhouw et al., 2010; Zhu et al., 2019). However, the capacity for these dually marked domains to persist during zebrafish gastrulation or for new domains of dual enrichment to emerge during this period has not been addressed.

For each histone modification, we performed two replicate CUT&RUN experiments using pools of 25 embryos per replicate. CUT&RUN was performed using an adapted version of the protocol first described by Skene et al for use in yeast and mammalian cells (Skene et al., 2018). A detailed version of this adapted protocol is included in supplementary materials. In total, we identified 29,562 sites of H3K4me3 enrichment and 9,948 sites of H3K27me3 enrichment. Consistent with previous studies, we observed strong enrichment of H3K4me3 within the 500 base pairs (bp) surrounding the TSS of known genes (**Figure 1A-C**). Enrichment of H3K27me3 was found at the TSS of a smaller subset of genes, with broad H3K27me3 peaks noted in additional regions including Hox gene clusters (**Figure 1A-D**). Consistent with known associations, published RNA-seq data from shield stage embryos (White et al., 2017) demonstrated the presence of RNA transcripts derived from 96% of genes exclusively marked by H3K4me3, while only 6% of those marked exclusively by H3K27me3 were associated with RNA transcripts at shield stage. Taken together, these data demonstrate that CUT&RUN can be used for robust detection of modified histones in the embryonic zebrafish genome.

**Figure 1:**
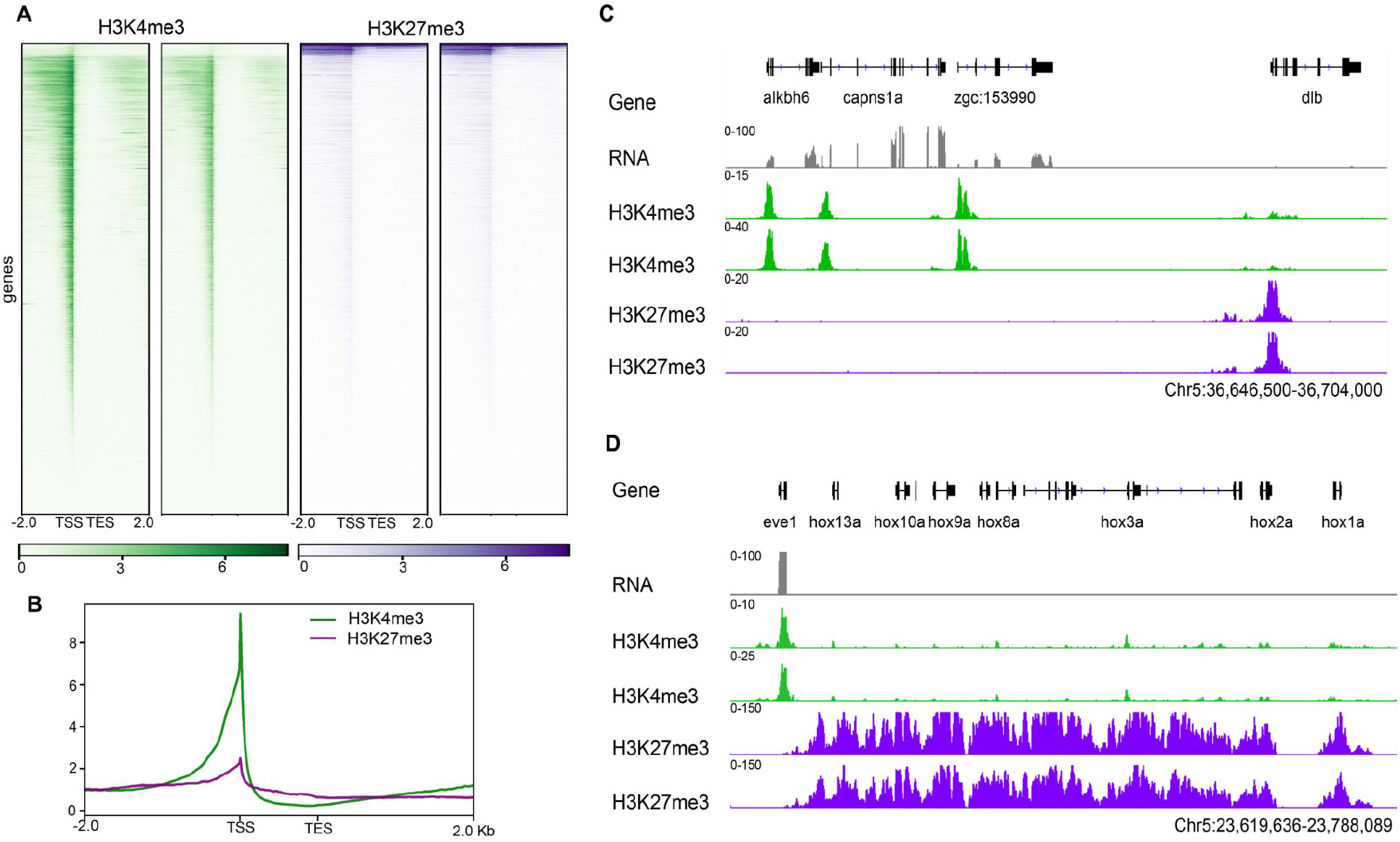
CUT&RUN detects H3K4me3 and H3K27me3 at the transcriptional start sites of annotated genes. **(A)** Heat maps of replicate data for H3K4me3 (green) and H3K27me3 (purple) enrichment as detected by CUT&RUN. (**B**) Profile plot of H3K4me3 and H3K27me3 at the TSS of annotated genes. **(C)** Genome browser views of H3K4me3 and H3K27me3 enrichment at select loci. H3K4me3 is detected at the TSS of three genes with associated RNA transcripts (*alkbh6, capns1a*, and *zgc153990*) and H3K27me3 is detected at the TSS of the inactive *dlb* gene. **(D)** Genome browser view of broad H3K27me3 enrichment over the hoxA gene cluster on chromosome 5. In **C** and **D** replicate data are shown as individual tracks.

Next, we investigated the co-occurrence of H3K4me3 and H3K27me3 in shield stage CUT&RUN data. In total, we detected 6,570 regions that were enriched for both H3K4me3 and H3K27me3 in shield stage embryos (**Figure 2A-B**). Notably, the majority of sites harboring both modifications in shield stage embryos (89%) do not show dual enrichment in previously published ChIP data from 1000-cell stage embryos (Zhu et al., 2019) (**Figure 2C**). Among regions enriched in both modified histones at shield stage, 58% of associated genes lacked RNA transcripts at the same stage, suggesting that these genes may be poised for future transcription. Consistent with this model, genes enriched in both H3K4me3 and H3K27me3 at their TSS in shield stage CUT&RUN data were heavily biased toward regulation of developmental processes (**Figure 2D**). Genes gaining strong bivalency during gastrulation have also been recently identified in mouse (Xiang et al., 2020), and there is significant overlap between dually marked genes in both systems. Analysis of CUT&RUN data revealed that 40% of genes acquiring strong bivalency during mouse gastrulation are also dually enriched with H3K4me3 and H3K27me3 in shield stage zebrafish embryos (**Supplementary Table 1**). Conservation of these dually marked regions during gastrulation suggests the potential for a conserved chromatin program involved in guiding cell fate decisions during germ layer formation.

**Table 1.** Antibodies used in this study.

| **Antibody** | **Catalog number** | **Dilution** |
| --- | --- | --- |
| H3K4me3 | ab8580 (abcam) | 1/100 |
| H3K27me3 | 07-449 (millipore) | 1/50 |
| H3K9me3 | ab8898 (abcam) | 1/100 |
| RNA pol-II<br>CTD (phospho S5) | ab5408 (abcam) | 1/75 |
| H2A.Z | 39113 (active motif) | 1/50 |
| IgG | ab46540 (abcam) | 1/100 |

**Figure 2.**
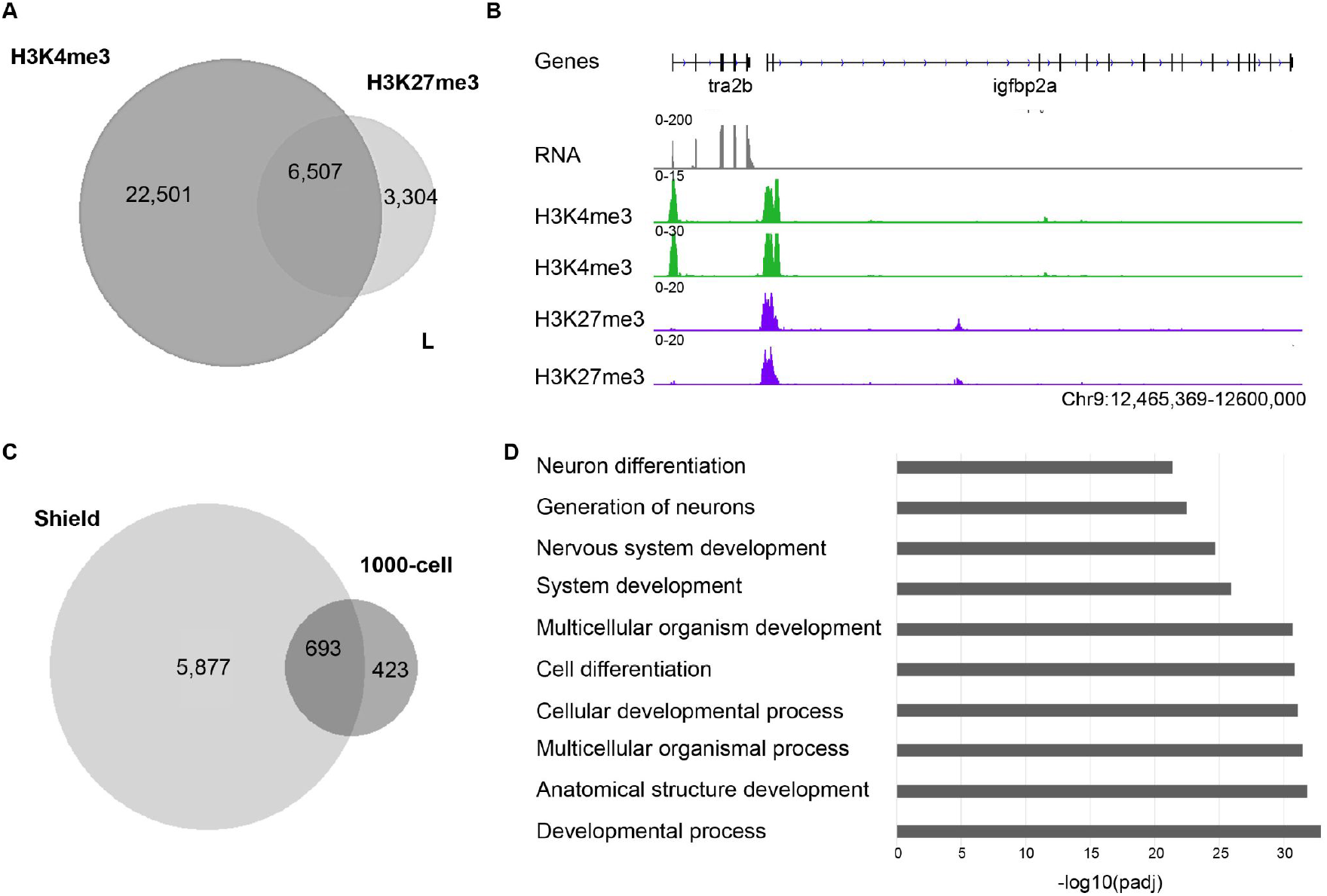
CUT&RUN identifies regions enriched in both H3K4me3 and H3K27me3 in shield stage zebrafish embryos. **(A)** Venn diagram indicating the number of regions enriched in H3K4me3, H3K27me3 or both modified histones in CUT&RUN data from shield stage embryos. Regions where one peak of H3K27me3 overlapped more than one peak of H3K4me3, or vice versa, were counted as one overlapping peak. **(B)** Genome browser view illustrating the co-occurrence of H3K4me3 and H3K27me3 enrichment peaks at the TSS of the *igfbp2a* gene, which lacks detectable transcripts at shield stage. At the same stage, the TSS of the nearby *tra2b* gene is exclusively marked by H3K4me3 and is associated with RNA transcripts. Replicate data are shown as individual tracks **(C)** Venn diagram showing shared and unique peaks of enrichment between CUT&RUN data from shield stage embryos and previously reported ChIP data from 1000-cell stage zebrafish embryos. **(D)** GO analysis reveals genes marked by both H3K4me3 and H3K27me3 at shield stage are associated with developmental processes including nervous system development.

### CUT&RUN effectively detects C-terminal domain serine 5 phosphorylated RNA pol II at the 5’ end of transcribed genes

To assess the potential for CUT&RUN to detect interactions between DNA and non-histone proteins, we performed CUT&RUN on shield stage embryos using an antibody that recognizes the serine 5 phosphorylated C-terminal domain of RNA polymerase II (Ser5P-pol II). This modified form of poI II is highly enriched near the TSS of genes undergoing active transcription (Chen et al., 2018; Harlen and Churchman, 2017). Consistent with known localization patterns, CUT&RUN revealed Ser5-pol II enrichment near the TSS of zebrafish genes, with 71% of enrichment peaks found at genes producing transcripts that were detectable by RNA-seq at shield stage (**Figure 3A-C**).

**Figure 3.**
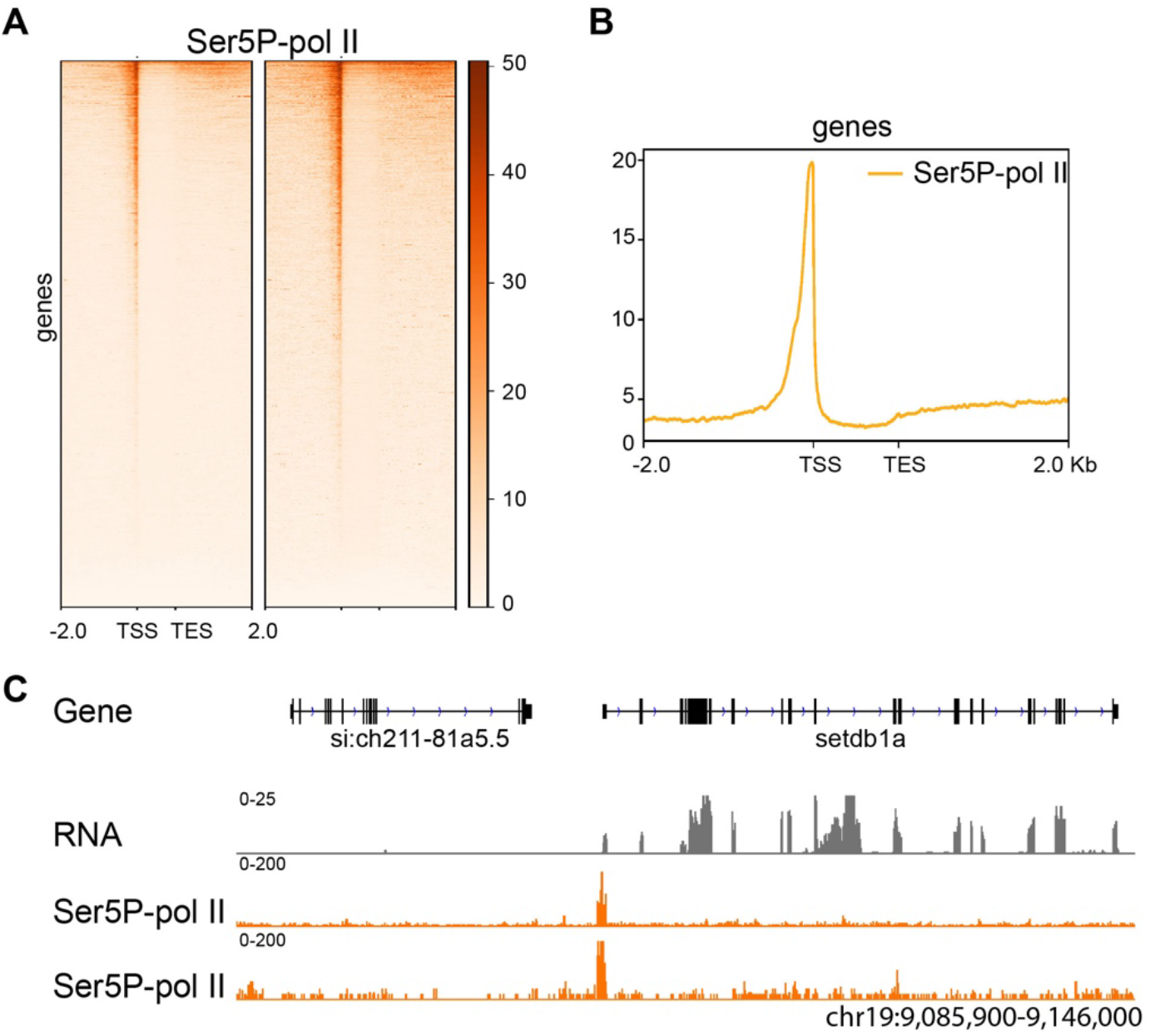
CUT&RUN detects Ser-5 CTD phosphorylated RNA polymerase 2 (Ser5P-pol II) at the TSS of transcribed genes. **(A)** Heat maps of replicate data for Ser5P-pol II enrichment at the TSS of annotated genes **(B)** Profile plot showing enrichment of Ser5-pol II at the TSS of annotated genes. **(C)** Genome browser view showing enrichment of Ser5-pol II at the start of *setdb1a*, which produces RNA transcripts at shield stage. Similar S5-pol II enrichment is not observed at the silent si:c*h211-81a5.5* gene. Replicate data are shown as individual tracks.

### CUT&RUN detects H3K9me3 enrichment at repetitive sequences, including in regions with low mappability

To assess CUT&RUN effectiveness in interrogating the repetitive fraction of the zebrafish genome, we examined enrichment of the heterochromatic histone modification H3K9me3 in shield stage embryos (**Figure 4A**). H3K9me3 is canonically associated with transcriptional silencing at repetitive sequences including many transposable elements (Allshire and Madhani, 2018; Janssen et al., 2018). As expected, only a small fraction (2.6%) of H3K9me3 enrichment peaks shared between replicates overlapped with transcriptional start sites of known genes, while 93% percent were associated with annotated repetitive sequences (**Figure 4B**). Among sites of enrichment, the largest number of peaks were associated with DNA transposons, followed by long terminal repeat (LTR) transposons and simple repeats (**Figure 4B**).

**Figure 4.**
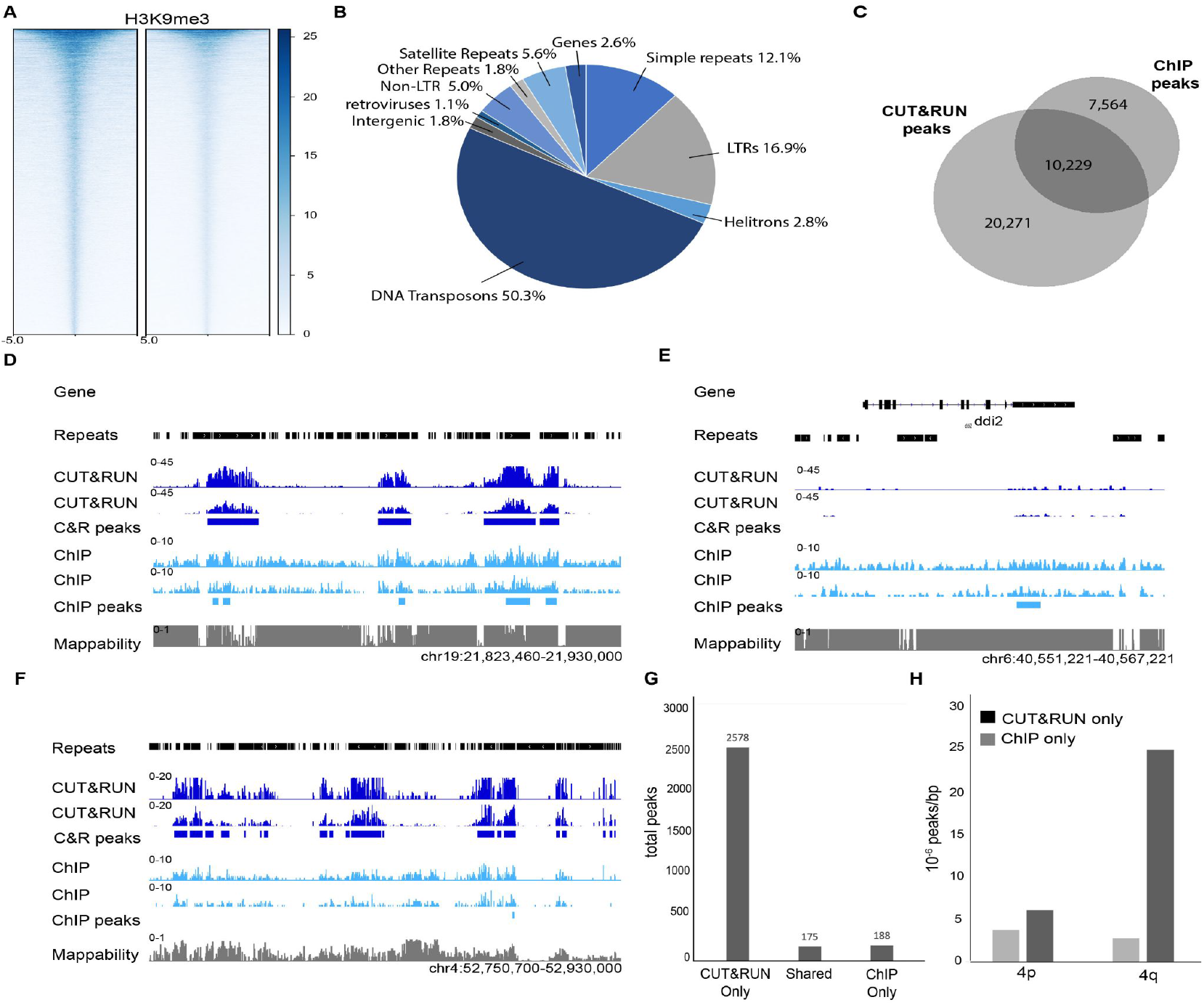
CUT&RUN detects H3K9me3 enrichment at repeated sequences. **(A)** Heat map of H3K9me3 enrichment around peak centers in CUT&RUN replicates **(B)** Pie chart depicting H3K9me3 enrichment at different repeat sequence classes as detected by CUT&RUN. **(C)** Venn diagram showing shared and unique peaks between shield stage CUT&RUN and ChIP data. **(D)** Genome browser view comparing enrichment and called peaks in CUT&RUN (dark blue) and ChIP (light blue) data from shield stage embryos. **(E)** Genome browser view of a select genomic loci containing a ChIP-only peak. **(F)** Genome browser view of a select genomic loci containing multiple CUT&RUN-only peaks on the long arm of chromosome 4. **(G)** Bar graph of CUT&RUN-only, shared and ChIP-only peaks detected in regions with mappability scores below 0.5 **(H)** Bar graph comparing CUT&RUN-only and ChIP-only peaks on the long vs short arms of chromosome 4. In **D-F** replicate data are shown as individual tracks.

We previously analyzed H3K9me3 in shield stage embryos by ChIP using the same antibody used for CUT&RUN analysis (Laue et al., 2019). Comparing H3K9me3 CUT&RUN data to published ChIP data revealed 10,229 sites of enrichment that were shared between ChIP and CUT&RUN datasets. We also noted 20,271 enrichment peaks that were unique to CUT&RUN data and 7,564 peaks were unique to ChIP (**Figure 4C**). Browser assessment of select loci suggested CUT&RUN yielded substantially improved signal to noise ratios over ChIP (**Figure 4D**), raising the possibility that some discrepancies could be due to improved accuracy in CUT&RUN peak calling. Consistent with this hypothesis, substantially improved Fraction of Reads in Peaks (FRiP) scores were observed for CUT&RUN data compared to ChIP (10.4% vs. 1.6%) (Landt et al., 2012), and browser views of select gene loci revealed ChIP-only peaks that were difficult to discriminate from surrounding background (**Figure 4E**).

In contrast to ChIP-only peaks, browser assessment of select CUT&RUN-only peaks revealed clear enrichment domains with signals well above background (**Figure 4F**). These peaks often appeared in repetitive, low-mappability regions where it can be difficult to map reads accurately (Pockrandt et al., 2020). Consistent with this initial inspection, we found that genome wide, there were roughly 10 times more CUT&RUN-only peaks in regions with mappability scores of less than 0.5 compared to ChIP-only or shared peaks (**Figure 4G**). This distinction was particularly noticeable on the long arm of chromosome 4 (4q), which is very highly enriched in repetitive elements and exists in a densely heterochromatic state (Anderson et al., 2012; Howe et al., 2013) (**Figure 4F, H**). Here, the number of CUT&RUN-only peaks was 4 times more frequent than on the euchromatic short arm (4p) of the same chromosome. In contrast, the number of ChIP-only peaks on 4p and 4q were similar. Collectively, these data suggest that the high signal-to-noise ratio and specificity of CUT&RUN make it especially well-suited for detecting protein-DNA interactions in repetitive regions.

### CUT&Tag effectively identifies sites enriched for the histone variant H2A.Z

Finally, we investigated whether CUT&Tag could also be applied for effective profiling of DNA-histone interactions in zebrafish. For this analysis, we examined the localization of the histone variant H2A.Z in shield stage embryos. In vertebrates, H2A.Z-containing nucleosomes are known to reside at gene promoters, and this histone variant has been implicated in protecting these sites from DNA methylation (Hu et al., 2013; Ku et al., 2012; Murphy et al., 2018). Previous studies have described H2A.Z localization in zebrafish sperm and blastula stage embryos by ChIP (Murphy et al., 2018). However, profiles have yet to be assessed during gastrulation or later stages of embryogenesis. CUT&Tag was performed using an adapted version of the protocol first described by Kaya-Okur, et al. for use in mammalian cells (Kaya-Okur et al., 2019). A detailed version of this adapted protocol is included in supplementary materials. Using this protocol, we detected robust signals for H2A.Z among replicates in shield stage embryos, and in keeping with previous findings, we found that H2A.Z was enriched at gene promoters (**Figure 5A-C**). This result demonstrates that CUT&Tag can be used effectively to detect H2A.Z enriched sites during zebrafish embryogenesis.

**Figure 5.**
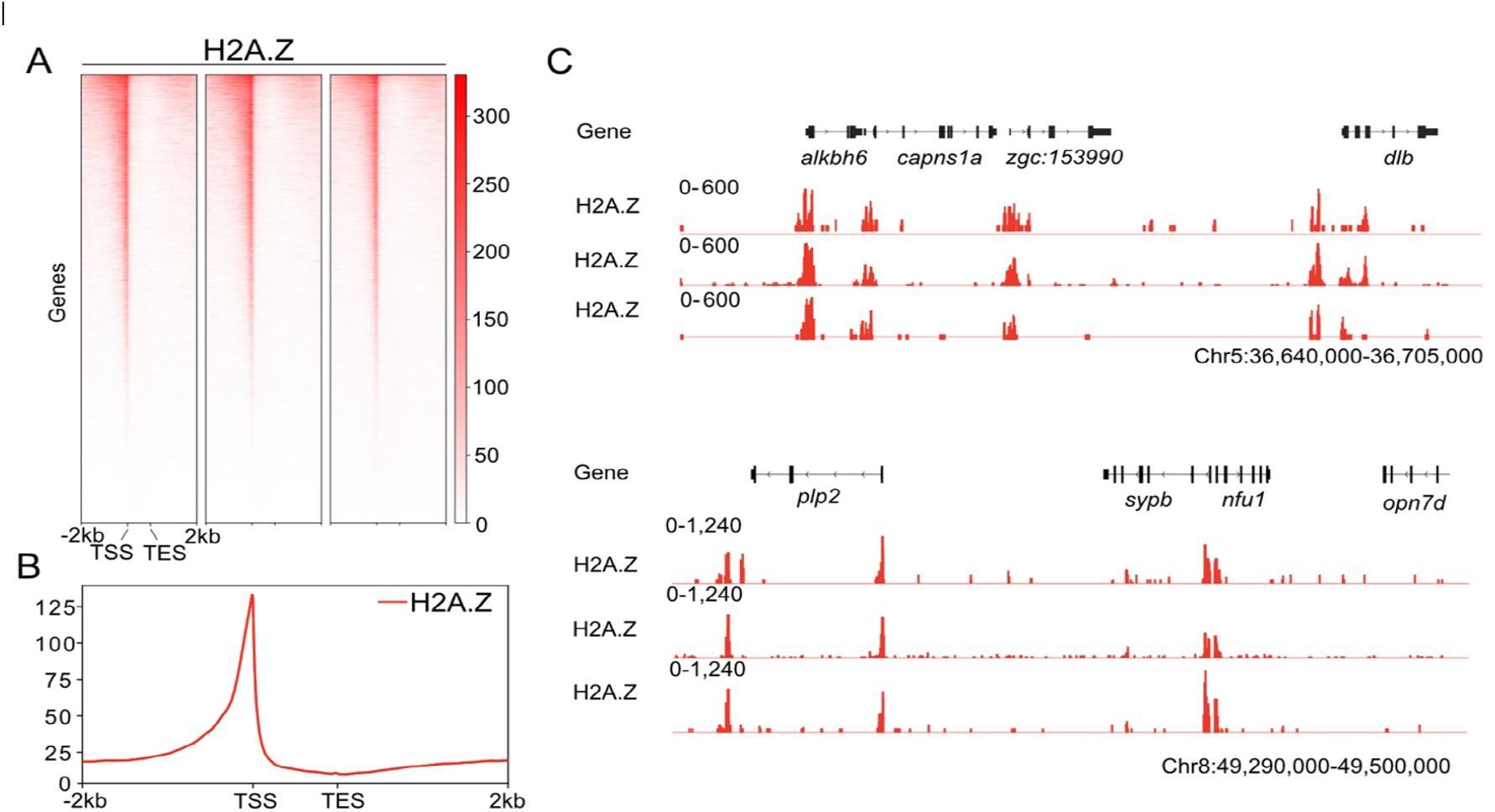
CUT&Tag detects H2A.Z in shield stage zebrafish embryos. **(A)** Heatmaps of replicate data for H2A.Z enrichment at annotated genes as detected by CUT&Tag in shield stage embryos. **(B)** Profile plot of H2A.Z at annotated genes. (C) Genome browser views of H2A.Z enrichment at selected loci. Replicate data for H2A.Z CUT&Tag are shown as individual track.

To further characterize potential changes in H2A.Z patterning during developmental transitions, we also performed CUT&Tag for H2A.Z using 24 hpf embryos. Correlation analysis showed that replicates were clustered according to developmental stages, indicating that H2A.Z enrichment patterns changed during developmental progression (**Figure 6A**). Further comparison of H2A.Z profiles between shield stage and 24 hpf embryos revealed that H2A.Z levels generally increased from shield stage to 24 hpf and that there were more sites that were unique to 24hpf embryos than to shield stage embryos (**Figure 6B-E**). Taken together these data suggest that H2A.Z profiles are dynamic and evolve over the course of embryogenesis.

**Figure 6.**
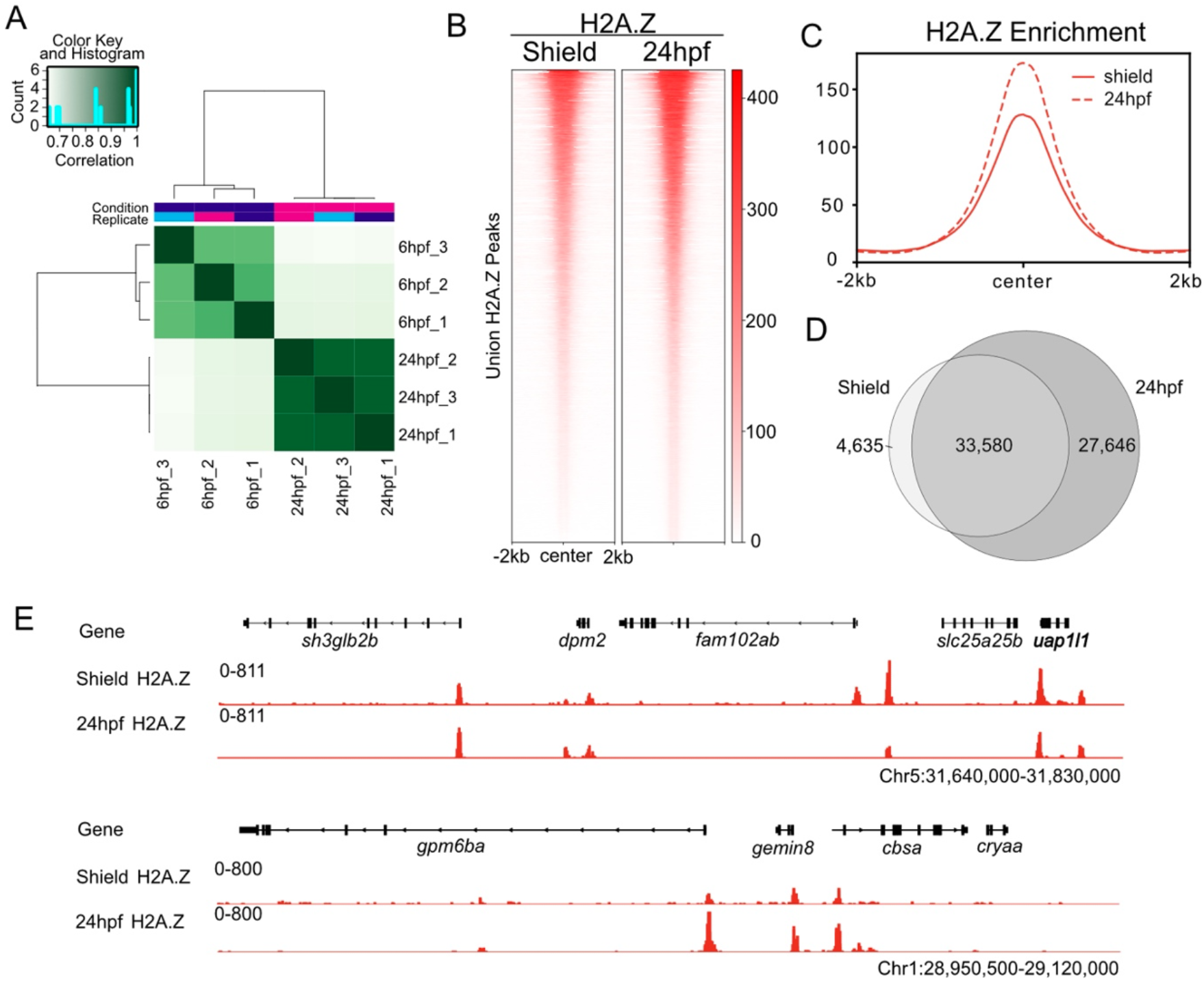
CUT&Tag identifies changes of H2A.Z patterning from shield stage to 24hpf zebrafish embryos. **(A) C**orrelation heatmap demonstrating high correlation among CUT&Tag replicates and separation based on developmental stages. **(B)** Heatmaps of H2A.Z enrichment at H2A.Z-marked regions as detected by CUT&Tag in shield stage and 24hpf embryos. Normalized H2A.Z signals were generated using merged replicate data. **(C)** Profile plot of H2A.Z enrichment at H2A.Z-marked regions in shield stage and 24hpf embryos. **(D)** Venn diagram showing shared and unique H2A.Z peaks between H2A.Z CUT&Tag data in shield stage and 24hpf embryos. (E) Genome browser views of H2A.Z enrichment in shield stage and 24hpf embryos at selected loci. Replicate data are shown as individual tracks.

## Discussion

Our results demonstrate the feasibility of using CUT&RUN and CUT&Tag approaches to profile chromatin during zebrafish embryogenesis, and provide the first high-resolution enrichment maps of H3K4me3, H3K27me3 and H3K9me3, Ser5P-pol II, and H2A.Z during zebrafish gastrulation. The reduced embryo requirements of CUT&RUN and CUT&Tag eliminate a significant barrier to current chromatin profiling during zebrafish embryogenesis, while increased signal-to-noise ratios and more precise foot printing afforded by CUT&RUN/CUT&Tag facilitate accurate identification of enrichment sites in these data sets.

In particular, our data demonstrate improved power of CUT&RUN for analysis of protein-DNA interactions that impact repetitive sequences. While we were initially surprised by the substantial number of non-overlapping peaks detected between H3K9me3 CUT&RUN and ChIP data, our analysis suggests that this discrepancy is due to improved enrichment peak calling at repetitive sequences, especially in regions of low mappability. Reduced background in CUT&RUN allows for effective global peak calling at lower sequencing depths than can be used for ChIP (Skene and Henikoff, 2017). This feature likely contributes to CUT&RUN’s increased effectiveness in low mappability regions, as more sequencing reads in these regions will typically be discarded due to low mapping quality.

It is important to appreciate that just like any antibody-based method, the specific efficacy of a given antibody-antigen interaction will be a critical determinant in whether CUT&RUN/CUT&Tag will be successful in profiling zebrafish chromatin. Given that these methods are performed on unfixed chromatin, in some cases, antibodies that perform well in other methods that lack fixation such as immunofluorescence may outperform those validated for ChIP. It is also important to note that higher salt concentrations needed for CUT&Tag make this method most suitable for assaying very stable interactions such as those between histones and DNA. In cases where interactions may be less stable, such as profiling transcription factor-DNA interactions, CUT&RUN is likely to be more effective (Kaya-Okur et al., 2019).

Our analysis of chromatin in shield stage zebrafish embryos provides an important reference for understanding how histone modifications and variants may help shape early cell fate decisions during development. For example, our analysis of shield stage CUT&RUN data suggests that many new bivalent domains emerge between the onset of zygotic transcription and zebrafish gastrulation. Although our data does not fully rule out the possibility that the dually marked domains detected specifically in shield stage zebrafish embryos result from mixed cell populations, or improved peak calling, overlap between domains identified in our data and those reported to gain strong bivalency during mouse gastrulation points toward the presence of a conserved chromatin program for directing early cell fate decisions.

Our data also reveal changes in H2A.Z deposition as embryos develop from shield stage to 24 hpf. A large number of enriched H2A.Z sites were called as peaks both at shield stage and at 24 hpf (88% of shield stage peaks), providing partial validation of our CUT&Tag methods, and suggesting that there is general maintenance of embryonic H2A.Z localization over this developmental period. In addition to these 33,580 maintained sites, *de novo* H2A.Z accumulated at 27,646 genomic locations in 24hpf embryos, indicating that broad expansion of H2A.Z localization occurs as embryos progress through segmentation stages of development. This shifting H2A.Z landscape suggests genomic H2A.Z reorganization may help drive differentiation over this developmental window. Consistent with this model, prior studies indicate that disruption of H2A.Z over similar developmental periods in zebrafish and frog embryos leads to defects in neural crest derived tissues (Greenberg et al., 2019; Raja et al., 2020).

Taken together, our data validate the use of CUT&RUN and CUT&Tag in zebrafish. The high-resolution chromatin maps of zebrafish embryos undergoing gastrulation generated in this study provide an important resource for probing the relationship between chromatin states and early cell fate decisions. At the same time, the detailed protocols for CUT&RUN and CUT&Tag accompanying this study will facilitate efficient adoption of these methods by the zebrafish field, allowing for large scale, high resolution, time course experiments to broadly define chromatin dynamics across the entirety of early zebrafish embryogenesis.

## Methods

### Zebrafish

Zebrafish husbandry and care were conducted in full accordance with animal care and use guidelines with ethical approval by the Institutional Animal Care and Use Committees at the University of Georgia and the University Committee on Animal Resources at the University of Rochester Medical Center. Zebrafish were raised and maintained under standard conditions in compliance with relevant protocols and ethical regulations. Fertilized eggs were obtained from the zebrafish AB strain. Embryos were reared in system water at 28.5 °C and staged according to morphology

### CUT&RUN and CUT&Tag

CUT&RUN and CUT&Tag were performed according to published methods, with minor adaptations (Kaya-Okur et al., 2019; Skene et al., 2018). Detailed protocols describing the exact CUT&RUN and CUT&Tag methods used in this study, and preparation of CUT&RUN libraries are provided in supplementary materials. Antibodies used in this study are provided in Table 1.

### Sequencing data

CUT&RUN libraries were pooled and sequenced on a NextSeq500 instrument at the Georgia Genomics Facility. Sequencing details are provided in Table 2, and raw data generated in this study can be found at NCBI GEO (accession GSE178343). CUT&Tag libraries from 24 hpf embryos were pooled and sequenced with services from NovoGene Co. on a HiSeq2500 instrument, and libraries from shield stage embryos were sequenced with Genewiz Co. on a NextSeq550 instrument. Sequencing details are provided in Table 2, and raw data generated in this study can be found at NCBI GEO (accession GSE178559). Publicly available RNA data used in this study can be found at the EBI European Nucleotide Archive (accession # PRJEB12982). Publicly available H3K4me3 & H3K27me3 ChIP data used in this study can be found at NCBI GEO (accession # GSE110663). Publicly available H3K9me3 ChIP data used in this study can be found at NCBI GEO (accession # GSE113086).

**Table 2:** Total raw, trimmed and aligned reads per sample.

| **Sample** | **Raw Reads** | **Trimmed Reads** | **Aligned Reads** | **% Aligned** |
| --- | --- | --- | --- | --- |
| CnR_K4_1 | 31,343,063 | 31,052,132 | 23,967,960 | 77.19% |
| CnR_K4_2 | 70,960,918 | 70,283,557 | 53,529,863 | 76.16% |
| CnR_K27_1 | 26,960,471 | 26,702,170 | 18,939,978 | 70.93% |
| CnR_K27_2 | 27,851,277 | 27,590,408 | 19,356,459 | 70.16% |
| CnR_K9_1 | 22,214,067 | 21,923,529 | 4,830,123 | 22.03% |
| CnR_K9_2 | 43,462,568 | 42,894,733 | 9,922,627 | 23.13% |
| CnR_pol2_1 | 9,842,850 | 9,831,381 | 7,214,488 | 73.38% |
| CnR_pol2_2 | 5,707,930 | 5,700,927 | 4,137,608 | 72.58% |
| CnR_IgG | 8,094,912 | 8,079,509 | 4,777,942 | 59.14% |
| CUT&Tag_6hpf_H2A.Z_1 | 1,233,216 | 1,233,216 | 1,126,474 | 91.34% |
| CUT&Tag_6hpf_H2A.Z_2 | 1,447,412 | 1,447,412 | 1,305,818 | 90.22% |
| CUT&Tag_6hpf_H2A.Z_3 | 3,135,635 | 3,135,635 | 2,847,270 | 90.80% |
| CUT&Tag_24hpf_H2A.Z_1 | 23,376,962 | 23,376,962 | 21,779,997 | 93.17% |
| CUT&Tag_24hpf_H2A.Z_2 | 28,652,092 | 28,652,092 | 26,947,558 | 94.05% |
| CUT&Tag_24hpf_H2A.Z_3 | 23,810,168 | 23,810,168 | 22,288,633 | 93.61% |

### CUT&RUN data analysis

Short reads (<20 bp) and adaptor sequences were removed using TrimGalore (version 0.6.5), cutadapt version 2.8, and Python 3.7.4, with fastqc command (version 0.11.9) (https://github.com/FelixKrueger/TrimGalore). Trimmed Illumina reads were aligned to the current zebrafish genome assembly (GRCz.11, Ensembl release 103) using Bowtie2 (version 2.4.1) with the “very-sensitive-local” parameters which assigns multi-mapped reads to their best alignment based on MAPQ scores (Langmead and Salzberg, 2012). Reads were also aligned to the spike-in genomes using the same parameters as above. For the histone modifications, the spike-in was *S. cerevisiae* (R64-1-1, Ensembl release 48) and for the pol-II, the spike-in was *E. coli* (assembly GCF_000005845.2_ASM584v2). Alignments were filtered using SAMtools (version 1.10) for a MAPQ score of 20 (Danecek et al., 2021; Li et al., 2009). Using a modified version of the script from Skene et al 2017, CUT&RUN data was normalized for spike-in and library-size with Bedtools “genomecov” (version 2.29.2) (Quinlan and Hall, 2010). This outputs a bedgraph file which was adapted to a bed file for peak-calling and converted to a bigwig using ucsc “bedGraphToBigWig” (version 359) for data visualization in the Integrated Genome Viewer. The Hypergeometric Optimization of Motif EnRichment (HOMER) software package (version 4.11) was used to identify peaks over input(Heinz et al., 2010). For pol-II CUT&RUN, the parameters used were -style factor -gsize 1.5e9. For histone modification CUT&RUN, the parameters used were -style histone -minDist 1000 -F 6 -gsize 1.5e9 -fdr 0.0001. Bedtools “intersect” was used to compare peak locations between samples (Quinlan and Hall, 2010). Peak annotation was performed using HOMER “annotatePeaks.pl” with the masked reference annotation (Heinz et al., 2010). Only peaks within 1000 bps of a TSS were considered to be associated with that gene. For the H3K9me3 CUT&RUN, peaks that were further than 1000 bps from a genic TSS were then re-annotated using an un-masked reference annotation to determine which enrichment domains were associated with repetitive elements. deepTools (version 3.3.1) was used to construct heatmaps (plotHeatmap) and metaplots (plotProfile) (Ramirez et al., 2016). Mappability was calculated with GenMap using a kmer of 76 bp and a mismatch allowance of 2 (Pockrandt et al., 2020). To identify H3K9me3 peaks associated with regions of low mappability, Bedtools “intersect” was used with -f 0.3, requiring at least 30% of a peak be covered with mappability scores <50%. Proportional venn diagrams were made with the help of BioVenn (Hulsen et al., 2008). Gene Ontology Analysis was performed using gProfiler (Raudvere et al., 2019). Scripts used for data processing can be found at https://github.com/klduval/CutNRun_2021.

### ChIP data analysis

Short reads (<20 bp) and adaptor sequences were removed using TrimGalore (version 0.6.5), cutadapt version 2.8, and Python 3.7.4, with fastqc command (version 0.11.9) (https://github.com/FelixKrueger/TrimGalore). Trimmed Illumina reads were aligned to the current zebrafish genome assembly (GRCz.11, Ensembl release 103) using Bowtie2 (version 2.4.1) with the “very-sensitive-local” parameters which assigns multi-mapped reads to their best alignment based on MAPQ scores. Zhu 2019 ChIP paired reads had become disordered so were repaired using BBMap “repair.sh” (version 38.83) before alignment. Alignments were filtered using SAMtools (version 1.10) for a MAPQ score of 20 (Danecek et al., 2021; Li et al., 2009). The Hypergeometric Optimization of Motif EnRichment (HOMER) software package (version 4.11) was used to identify peaks over input with the following parameters: -style histone-size 1000 -gsize 1.5e9. Bedtools “intersect” was used to compare peak locations between samples (Quinlan and Hall, 2010). To plot the relative distribution of mapped ChIP reads, read counts were determined for each 10 bp window across the genome using deepTools “bamCoverage” (version 3.3.1) and data were displayed using the Integrated Genome Viewer (Ramirez et al., 2016).

### CUT&Tag data analysis

For H2A.Z CUT&Tag paired-end sequencing reads, adaptor sequences were removed using cutadapt (version 2.7). Trimmed reads were aligned to zebrafish genome assembly (GRCz11) using Bowtie2 (version 2.2.5) with default parameters (Langmead and Salzberg, 2012). Unmapped reads were filtered using samtools (version 1.9), and PCR duplicates were removed using picard MarkDuplicates (version 2.5.0) (Danecek et al., 2021; Li et al., 2009). H2A.Z replicate data were merged using picard MergeSamFiles, and genome browser tracks were generated using DeepTOOLS bamCoverage (version 3.5.1) with the following setting: -- normalizeUsing RPKM --binSize 10 (Ramirez et al., 2016). Peak calling was performed using macs2 callpeak (2.2.6) with the following parameters: -f BAMPE -g 1.5e9 --nomodel --broad. Heatmaps and profile plots were generated using plotHeatmap and plotProfile from deeptools (version 3.5.1), and fonts and labels were adjusted in Affinity Designer (version 1.9.1) (Ramirez et al., 2016). Overlapping peak analysis was performed using bedtools intersect (version 2.30.0), and venn diagram was generated using R package eulerr (version 6.1.0 and R version 4.0.3) (Larsson, 2020)

### RNA-seq data analysis

Short reads (<20 bp) and adaptor sequences were removed using TrimGalore (version 0.6.5), cutadapt version 2.8, and Python 3.7.4, with fastqc command (version 0.11.9). Trimmed Illumina reads were aligned to the current zebrafish genome assembly (GRCz.11, Ensembl release 103) using STAR (version 2.7.3a) with option “--outSAMmultNmax 1” to keep multimapping reads but only report their best alignment (Dobin et al., 2013). TPMCalculator (version 0.0.4) was used to determine which genes are actively expressed (TPM >0.5) at the shield stage (Vera Alvarez et al., 2019). For pol-II CUT&RUN, Bedtools “intersect” was used to compare peaks with expressed genes. For H3K4me3 and H3K27me3 CUT&RUN, Bedtools “intersect” was also used to compare peaks within 1000bps of a TSS with expressed exons to ascertain which domains are associated with active transcription.

## Supporting information

Detailed CUT&RUN and CUT&Tag Protocols

## Acknowledgements

This work was supported by the National Institutes of Health (R01GM110092) to M.G.G, (T32GM007103) to K.L.D and (R35GM137833) to P.J.M. We thank Bob Schmitz and Pablo Mendieta for helpful advice on data analysis, and Felicia Ebot-Ojong and Zack Lewis for providing purified pA/GMnase protein. Sequencing of CUT&RUN samples was performed by the Georgia Genomics and Bioinformatics Core at the University of Georgia. Content is solely the responsibility of the authors and does not necessarily represent official views of the National Institutes of Health.

## Supplemental information

**Supplementary Table 1:** List of genes with super bivalency during mouse gastrulation, that are also marked by both H3K4me3 and H3K27me3 in shield stage zebrafish embryos.

## References

Ahmad, K., Spens, A.E., 2019. Separate Polycomb Response Elements control chromatin state and activation of the vestigial gene. PLoS Genet 15, e1007877.

Akdogan-Ozdilek, B., Duval, K.L., Goll, M.G., 2020. Chromatin dynamics at the maternal to zygotic transition: recent advances from the zebrafish model. F1000Res 9.

Allshire, R.C., Madhani, H.D., 2018. Ten principles of heterochromatin formation and function. Nat Rev Mol Cell Biol 19, 229–244.

Anderson, J.L., Rodriguez Mari, A., Braasch, I., Amores, A., Hohenlohe, P., Batzel, P., Postlethwait, J.H., 2012. Multiple sex-associated regions and a putative sex chromosome in zebrafish revealed by RAD mapping and population genomics. PLoS One 7, e40701.

Barski, A., Cuddapah, S., Cui, K., Roh, T.Y., Schones, D.E., Wang, Z., Wei, G., Chepelev, I., Zhao, K., 2007. High-resolution profiling of histone methylations in the human genome. Cell 129, 823–837.

Bartosovic, M., Kabbe, M., Castelo-Branco, G., 2021. Single-cell CUT&Tag profiles histone modifications and transcription factors in complex tissues. Nat Biotechnol.

Bernstein, B.E., Mikkelsen, T.S., Xie, X., Kamal, M., Huebert, D.J., Cuff, J., Fry, B., Meissner, A., Wernig, M., Plath, K., Jaenisch, R., Wagschal, A., Feil, R., Schreiber, S.L., Lander, E.S., 2006. A bivalent chromatin structure marks key developmental genes in embryonic stem cells. Cell 125, 315–326.

Brind’Amour, J., Liu, S., Hudson, M., Chen, C., Karimi, M.M., Lorincz, M.C., 2015. An ultra-low-input native ChIP-seq protocol for genome-wide profiling of rare cell populations. Nat Commun 6, 6033.

Chen, F.X., Smith, E.R., Shilatifard, A., 2018. Born to run: control of transcription elongation by RNA polymerase II. Nat Rev Mol Cell Biol 19, 464–478.

Danecek, P., Bonfield, J.K., Liddle, J., Marshall, J., Ohan, V., Pollard, M.O., Whitwham, A., Keane, T., McCarthy, S.A., Davies, R.M., Li, H., 2021. Twelve years of SAMtools and BCFtools. Gigascience 10.

Daneshvar, K., Ardehali, M.B., Klein, I.A., Hsieh, F.K., Kratkiewicz, A.J., Mahpour, A., Cancelliere, S.O.L., Zhou, C., Cook, B.M., Li, W., Pondick, J.V., Gupta, S.K., Moran, S.P., Young, R.A., Kingston, R.E., Mullen, A.C., 2020. lncRNA DIGIT and BRD3 protein form phase-separated condensates to regulate endoderm differentiation. Nat Cell Biol 22, 1211–1222.

Dobin, A., Davis, C.A., Schlesinger, F., Drenkow, J., Zaleski, C., Jha, S., Batut, P., Chaisson, M., Gingeras, T.R., 2013. STAR: ultrafast universal RNA-seq aligner. Bioinformatics 29, 15–21.

Ernst, C., Eling, N., Martinez-Jimenez, C.P., Marioni, J.C., Odom, D.T., 2019. Staged developmental mapping and X chromosome transcriptional dynamics during mouse spermatogenesis. Nat Commun 10, 1251.

Federation, A.J., Nandakumar, V., Searle, B.C., Stergachis, A., Wang, H., Pino, L.K., Merrihew, G., Ting, Y.S., Howard, N., Kutyavin, T., MacCoss, M.J., Stamatoyannopoulos, J.A., 2020. Highly Parallel Quantification and Compartment Localization of Transcription Factors and Nuclear Proteins. Cell Rep 30, 2463–2471 e2465.

Greenberg, R.S., Long, H.K., Swigut, T., Wysocka, J., 2019. Single Amino Acid Change Underlies Distinct Roles of H2A.Z Subtypes in Human Syndrome. Cell 178, 1421–1436 e1424.

Hainer, S.J., Fazzio, T.G., 2019. High-Resolution Chromatin Profiling Using CUT&RUN. Curr Protoc Mol Biol 126, e85.

Harlen, K.M., Churchman, L.S., 2017. The code and beyond: transcription regulation by the RNA polymerase II carboxy-terminal domain. Nat Rev Mol Cell Biol 18, 263–273.

Heinz, S., Benner, C., Spann, N., Bertolino, E., Lin, Y.C., Laslo, P., Cheng, J.X., Murre, C., Singh, H., Glass, C.K., 2010. Simple combinations of lineage-determining transcription factors prime cis-regulatory elements required for macrophage and B cell identities. Mol Cell 38, 576–589.

Horsfield, J.A., 2019. Packaging development: how chromatin controls transcription in zebrafish embryogenesis. Biochem Soc Trans 47, 713–724.

Howe, K., Clark, M.D., Torroja, C.F., Torrance, J., Berthelot, C., Muffato, M., Collins, J.E., Humphray, S., McLaren, K., Matthews, L., McLaren, S., Sealy, I., Caccamo, M., Churcher, C., Scott, C., Barrett, J.C., Koch, R., Rauch, G.J., White, S., Chow, W., Kilian, B., Quintais, L.T., Guerra-Assuncao, J.A., Zhou, Y., Gu, Y., Yen, J., Vogel, J.H., Eyre, T., Redmond, S., Banerjee, R., Chi, J., Fu, B., Langley, E., Maguire, S.F., Laird, G.K., Lloyd, D., Kenyon, E., Donaldson, S., Sehra, H., Almeida-King, J., Loveland, J., Trevanion, S., Jones, M., Quail, M., Willey, D., Hunt, A., Burton, J., Sims, S., McLay, K., Plumb, B., Davis, J., Clee, C., Oliver, K., Clark, R., Riddle, C., Elliot, D., Threadgold, G., Harden, G., Ware, D., Begum, S., Mortimore, B., Kerry, G., Heath, P., Phillimore, B., Tracey, A., Corby, N., Dunn, M., Johnson, C., Wood, J., Clark, S., Pelan, S., Griffiths, G., Smith, M., Glithero, R., Howden, P., Barker, N., Lloyd, C., Stevens, C., Harley, J., Holt, K., Panagiotidis, G., Lovell, J., Beasley, H., Henderson, C., Gordon, D., Auger, K., Wright, D., Collins, J., Raisen, C., Dyer, L., Leung, K., Robertson, L., Ambridge, K., Leongamornlert, D., McGuire, S., Gilderthorp, R., Griffiths, C., Manthravadi, D., Nichol, S., Barker, G., Whitehead, S., Kay, M., Brown, J., Murnane, C., Gray, E., Humphries, M., Sycamore, N., Barker, D., Saunders, D., Wallis, J., Babbage, A., Hammond, S., Mashreghi-Mohammadi, M., Barr, L., Martin, S., Wray, P., Ellington, A., Matthews, N., Ellwood, M., Woodmansey, R., Clark, G., Cooper, J., Tromans, A., Grafham, D., Skuce, C., Pandian, R., Andrews, R., Harrison, E., Kimberley, A., Garnett, J., Fosker, N., Hall, R., Garner, P., Kelly, D., Bird, C., Palmer, S., Gehring, I., Berger, A., Dooley, C.M., Ersan-Urun, Z., Eser, C., Geiger, H., Geisler, M., Karotki, L., Kirn, A., Konantz, J., Konantz, M., Oberlander, M., Rudolph-Geiger, S., Teucke, M., Lanz, C., Raddatz, G., Osoegawa, K., Zhu, B., Rapp, A., Widaa, S., Langford, C., Yang, F., Schuster, S.C., Carter, N.P., Harrow, J., Ning, Z., Herrero, J., Searle, S.M., Enright, A., Geisler, R., Plasterk, R.H., Lee, C., Westerfield, M., de Jong, P.J., Zon, L.I., Postlethwait, J.H., Nusslein-Volhard, C., Hubbard, T.J., Roest Crollius, H., Rogers, J., Stemple, D.L., 2013. The zebrafish reference genome sequence and its relationship to the human genome. Nature 496, 498–503.

Hu, G., Cui, K., Northrup, D., Liu, C., Wang, C., Tang, Q., Ge, K., Levens, D., Crane-Robinson, C., Zhao, K., 2013. H2A.Z facilitates access of active and repressive complexes to chromatin in embryonic stem cell self-renewal and differentiation. Cell Stem Cell 12, 180–192.

Hulsen, T., de Vlieg, J., Alkema, W., 2008. BioVenn - a web application for the comparison and visualization of biological lists using area-proportional Venn diagrams. BMC Genomics 9, 488.

Hyle, J., Zhang, Y., Wright, S., Xu, B., Shao, Y., Easton, J., Tian, L., Feng, R., Xu, P., Li, C., 2019. Acute depletion of CTCF directly affects MYC regulation through loss of enhancer-promoter looping. Nucleic Acids Res 47, 6699–6713.

Inoue, A., Chen, Z., Yin, Q., Zhang, Y., 2018. Maternal Eed knockout causes loss of H3K27me3 imprinting and random X inactivation in the extraembryonic cells. Genes Dev 32, 1525–1536.

Janssen, A., Colmenares, S.U., Karpen, G.H., 2018. Heterochromatin: Guardian of the Genome. Annu Rev Cell Dev Biol 34, 265–288.

Johnson, D.S., Mortazavi, A., Myers, R.M., Wold, B., 2007. Genome-wide mapping of in vivo protein-DNA interactions. Science 316, 1497–1502.

Kasinathan, S., Orsi, G.A., Zentner, G.E., Ahmad, K., Henikoff, S., 2014. High-resolution mapping of transcription factor binding sites on native chromatin. Nat Methods 11, 203–209.

Kaya-Okur, H.S., Wu, S.J., Codomo, C.A., Pledger, E.S., Bryson, T.D., Henikoff, J.G., Ahmad, K., Henikoff, S., 2019. CUT&Tag for efficient epigenomic profiling of small samples and single cells. Nat Commun 10, 1930.

Kong, N.R., Bassal, M.A., Tan, H.K., Kurland, J.V., Yong, K.J., Young, J.J., Yang, Y., Li, F., Lee, J.D., Liu, Y., Wu, C.S., Stein, A., Luo, H.R., Silberstein, L.E., Bulyk, M.L., Tenen, D.G., Chai, L., 2021. Zinc Finger Protein SALL4 Functions through an AT-Rich Motif to Regulate Gene Expression. Cell Rep 34, 108574.

Ku, M., Jaffe, J.D., Koche, R.P., Rheinbay, E., Endoh, M., Koseki, H., Carr, S.A., Bernstein, B.E., 2012. H2A.Z landscapes and dual modifications in pluripotent and multipotent stem cells underlie complex genome regulatory functions. Genome Biol 13, R85.

Landt, S.G., Marinov, G.K., Kundaje, A., Kheradpour, P., Pauli, F., Batzoglou, S., Bernstein, B.E., Bickel, P., Brown, J.B., Cayting, P., Chen, Y., DeSalvo, G., Epstein, C., Fisher-Aylor, K.I., Euskirchen, G., Gerstein, M., Gertz, J., Hartemink, A.J., Hoffman, M.M., Iyer, V.R., Jung, Y.L., Karmakar, S., Kellis, M., Kharchenko, P.V., Li, Q., Liu, T., Liu, X.S., Ma, L., Milosavljevic, A., Myers, R.M., Park, P.J., Pazin, M.J., Perry, M.D., Raha, D., Reddy, T.E., Rozowsky, J., Shoresh, N., Sidow, A., Slattery, M., Stamatoyannopoulos, J.A., Tolstorukov, M.Y., White, K.P., Xi, S., Farnham, P.J., Lieb, J.D., Wold, B.J., Snyder, M., 2012. ChIP-seq guidelines and practices of the ENCODE and modENCODE consortia. Genome Res 22, 1813–1831.

Langmead, B., Salzberg, S.L., 2012. Fast gapped-read alignment with Bowtie 2. Nat Methods 9, 357–359.

Larsson, J., 2020. eulerr:Area-Proportional Euler and Venn Diagrams and Ellipses. R package version.

Laue, K., Rajshekar, S., Courtney, A.J., Lewis, Z.A., Goll, M.G., 2019. The maternal to zygotic transition regulates genome-wide heterochromatin establishment in the zebrafish embryo. Nat Commun 10, 1551.

Lawrence, M., Daujat, S., Schneider, R., 2016. Lateral Thinking: How Histone Modifications Regulate Gene Expression. Trends Genet 32, 42–56.

Li, H., Handsaker, B., Wysoker, A., Fennell, T., Ruan, J., Homer, N., Marth, G., Abecasis, G., Durbin, R., Genome Project Data Processing, S., 2009. The Sequence Alignment/Map format and SAMtools. Bioinformatics 25, 2078–2079.

Lindeman, L.C., Andersen, I.S., Reiner, A.H., Li, N., Aanes, H., Ostrup, O., Winata, C., Mathavan, S., Muller, F., Alestrom, P., Collas, P., 2011. Prepatterning of developmental gene expression by modified histones before zygotic genome activation. Dev Cell 21, 993–1004.

Liu, N., Hargreaves, V.V., Zhu, Q., Kurland, J.V., Hong, J., Kim, W., Sher, F., Macias-Trevino, C., Rogers, J.M., Kurita, R., Nakamura, Y., Yuan, G.C., Bauer, D.E., Xu, J., Bulyk, M.L., Orkin, S.H., 2018. Direct Promoter Repression by BCL11A Controls the Fetal to Adult Hemoglobin Switch. Cell 173, 430–442 e417.

Meers, M.P., Bryson, T.D., Henikoff, J.G., Henikoff, S., 2019. Improved CUT&RUN chromatin profiling tools. Elife 8.

Menon, D.U., Shibata, Y., Mu, W., Magnuson, T., 2019. Mammalian SWI/SNF collaborates with a polycomb-associated protein to regulate male germline transcription in the mouse. Development 146.

Murphy, K.E., Meng, F.W., Makowski, C.E., Murphy, P.J., 2020. Genome-wide chromatin accessibility is restricted by ANP32E. Nat Commun 11, 5063.

Murphy, P.J., Wu, S.F., James, C.R., Wike, C.L., Cairns, B.R., 2018. Placeholder Nucleosomes Underlie Germline-to-Embryo DNA Methylation Reprogramming. Cell 172, 993–1006 e1013.

Oomen, M.E., Hansen, A.S., Liu, Y., Darzacq, X., Dekker, J., 2019. CTCF sites display cell cycle-dependent dynamics in factor binding and nucleosome positioning. Genome Res 29, 236–249.

Pagin, M., Pernebrink, M., Giubbolini, S., Barone, C., Sambruni, G., Zhu, Y., Chiara, M., Ottolenghi, S., Pavesi, G., Wei, C.L., Cantu, C., Nicolis, S.K., 2021. Sox2 controls neural stem cell self-renewal through a Fos-centered gene regulatory network. Stem Cells.

Park, P.J., 2009. ChIP-seq: advantages and challenges of a maturing technology. Nat Rev Genet 10, 669–680.

Park, S.M., Cho, H., Thornton, A.M., Barlowe, T.S., Chou, T., Chhangawala, S., Fairchild, L., Taggart, J., Chow, A., Schurer, A., Gruet, A., Witkin, M.D., Kim, J.H., Shevach, E.M., Krivtsov, A., Armstrong, S.A., Leslie, C., Kharas, M.G., 2019. IKZF2 Drives Leukemia Stem Cell Self-Renewal and Inhibits Myeloid Differentiation. Cell Stem Cell 24, 153–165 e157.

Pockrandt, C., Alzamel, M., Iliopoulos, C.S., Reinert, K., 2020. GenMap: ultra-fast computation of genome mappability. Bioinformatics 36, 3687–3692.

Prakash, S.A., Barau, J., 2021. Chromatin Profiling in Mouse Embryonic Germ Cells by CUT&RUN. Methods Mol Biol 2214, 253–264.

Pritykin, Y., van der Veeken, J., Pine, A.R., Zhong, Y., Sahin, M., Mazutis, L., Pe’er, D., Rudensky, A.Y., Leslie, C.S., 2021. A unified atlas of CD8 T cell dysfunctional states in cancer and infection. Mol Cell 81, 2477–2493 e2410.

Quinlan, A.R., Hall, I.M., 2010. BEDTools: a flexible suite of utilities for comparing genomic features. Bioinformatics 26, 841–842.

Raja, D.A., Subramaniam, Y., Aggarwal, A., Gotherwal, V., Babu, A., Tanwar, J., Motiani, R.K., Sivasubbu, S., Gokhale, R.S., Natarajan, V.T., 2020. Histone variant dictates fate biasing of neural crest cells to melanocyte lineage. Development 147.

Ramirez, F., Ryan, D.P., Gruning, B., Bhardwaj, V., Kilpert, F., Richter, A.S., Heyne, S., Dundar, F., Manke, T., 2016. deepTools2: a next generation web server for deep-sequencing data analysis. Nucleic Acids Res 44, W160–165.

Raudvere, U., Kolberg, L., Kuzmin, I., Arak, T., Adler, P., Peterson, H., Vilo, J., 2019. g:Profiler: a web server for functional enrichment analysis and conversions of gene lists (2019 update). Nucleic Acids Res 47, W191–W198.

Roth, T.L., Puig-Saus, C., Yu, R., Shifrut, E., Carnevale, J., Li, P.J., Hiatt, J., Saco, J., Krystofinski, P., Li, H., Tobin, V., Nguyen, D.N., Lee, M.R., Putnam, A.L., Ferris, A.L., Chen, J.W., Schickel, J.N., Pellerin, L., Carmody, D., Alkorta-Aranburu, G., Del Gaudio, D., Matsumoto, H., Morell, M., Mao, Y., Cho, M., Quadros, R.M., Gurumurthy, C.B., Smith, B., Haugwitz, M., Hughes, S.H., Weissman, J.S., Schumann, K., Esensten, J.H., May, A.P., Ashworth, A., Kupfer, G.M., Greeley, S.A.W., Bacchetta, R., Meffre, E., Roncarolo, M.G., Romberg, N., Herold, K.C., Ribas, A., Leonetti, M.D., Marson, A., 2018. Reprogramming human T cell function and specificity with non-viral genome targeting. Nature 559, 405–409.

Schmid, M., Durussel, T., Laemmli, U.K., 2004. ChIC and ChEC; genomic mapping of chromatin proteins. Mol Cell 16, 147–157.

Schulz, K.N., Harrison, M.M., 2019. Mechanisms regulating zygotic genome activation. Nat Rev Genet 20, 221–234.

Skene, P.J., Henikoff, J.G., Henikoff, S., 2018. Targeted in situ genome-wide profiling with high efficiency for low cell numbers. Nat Protoc 13, 1006–1019.

Skene, P.J., Henikoff, S., 2017. An efficient targeted nuclease strategy for high-resolution mapping of DNA binding sites. Elife 6.

Solomon, M.J., Varshavsky, A., 1985. Formaldehyde-mediated DNA-protein crosslinking: a probe for in vivo chromatin structures. Proc Natl Acad Sci U S A 82, 6470–6474.

Talbert, P.B., Meers, M.P., Henikoff, S., 2019. Old cogs, new tricks: the evolution of gene expression in a chromatin context. Nat Rev Genet 20, 283–297.

Tao, Y., Liu, J., Guan, X., Chen, H., Ren, X., Wang, S., Ji, M., 2020. Estimation of potential agricultural non-point source pollution for Baiyangdian Basin, China, under different environment protection policies. PLoS One 15, e0239006.

Uyehara, C.M., McKay, D.J., 2019. Direct and widespread role for the nuclear receptor EcR in mediating the response to ecdysone in Drosophila. Proc Natl Acad Sci U S A 116, 9893–9902.

Vastenhouw, N.L., Zhang, Y., Woods, I.G., Imam, F., Regev, A., Liu, X.S., Rinn, J., Schier, A.F., 2010. Chromatin signature of embryonic pluripotency is established during genome activation. Nature 464, 922–926.

Vera Alvarez, R., Pongor, L.S., Marino-Ramirez, L., Landsman, D., 2019. TPMCalculator: one-step software to quantify mRNA abundance of genomic features. Bioinformatics 35, 1960–1962.

Wang, K., Wang, H., Li, C., Yin, Z., Xiao, R., Li, Q., Xiang, Y., Wang, W., Huang, J., Chen, L., Fang, P., Liang, K., 2021. Genomic profiling of native R loops with a DNA-RNA hybrid recognition sensor. Sci Adv 7.

White, R.J., Collins, J.E., Sealy, I.M., Wali, N., Dooley, C.M., Digby, Z., Stemple, D.L., Murphy, D.N., Billis, K., Hourlier, T., Fullgrabe, A., Davis, M.P., Enright, A.J., Busch-Nentwich, E.M., 2017. A high-resolution mRNA expression time course of embryonic development in zebrafish. Elife 6.

Wu, E., Vastenhouw, N.L., 2020. From mother to embryo: A molecular perspective on zygotic genome activation. Curr Top Dev Biol 140, 209–254.

Wu, S.J., Furlan, S.N., Mihalas, A.B., Kaya-Okur, H.S., Feroze, A.H., Emerson, S.N., Zheng, Y., Carson, K., Cimino, P.J., Keene, C.D., Sarthy, J.F., Gottardo, R., Ahmad, K., Henikoff, S., Patel, A.P., 2021. Single-cell CUT&Tag analysis of chromatin modifications in differentiation and tumor progression. Nat Biotechnol.

Xiang, Y., Zhang, Y., Xu, Q., Zhou, C., Liu, B., Du, Z., Zhang, K., Zhang, B., Wang, X., Gayen, S., Liu, L., Wang, Y., Li, Y., Wang, Q., Kalantry, S., Li, L., Xie, W., 2020. Epigenomic analysis of gastrulation identifies a unique chromatin state for primed pluripotency. Nat Genet 52, 95–105.

Xu, R., Li, C., Liu, X., Gao, S., 2021. Insights into epigenetic patterns in mammalian early embryos. Protein Cell 12, 7–28.

Yu, S., Guo, J., Sun, Z., Lin, C., Tao, H., Zhang, Q., Cui, Y., Zuo, H., Lin, Y., Chen, S., Liu, H., Chen, Z., 2021. BMP2-dependent gene regulatory network analysis reveals Klf4 as a novel transcription factor of osteoblast differentiation. Cell Death Dis 12, 197.

Zhang, W., Chen, Z., Yin, Q., Zhang, D., Racowsky, C., Zhang, Y., 2019. Maternal-biased H3K27me3 correlates with paternal-specific gene expression in the human morula. Genes Dev 33, 382–387.

Zheng, X.Y., Gehring, M., 2019. Low-input chromatin profiling in Arabidopsis endosperm using CUT&RUN. Plant Reprod 32, 63–75.

Zhu, W., Xu, X., Wang, X., Liu, J., 2019. Reprogramming histone modification patterns to coordinate gene expression in early zebrafish embryos. BMC Genomics 20, 248.

