## Supplementary material for "Identification of chromatin states during zebrafish gastrulation using CUT&RUN and CUT&Tag": Detailed CUT&RUN and CUT&Tag Protocols

### Additional Supplementary Information

#### I: Protocol for CUT&RUN analysis of 6hpf zebrafish embryos

##### A. DNA isolation:

The experimental protocol was adapted from Skene *et. al.*, Nature protocols, 2018,

The protocol is optimized for analysis of 6 hpf zebrafish embryos

**Key reagents:**

Concanavalin-A coated magnetic beads (Bangs laboratories: cat. No. BP531)

Digitonin (EMD Millipore: cat. No. 300410)

pAG-MNase, 20X stock (EpiCypher: 15-1116, or made in house)

Halt Protease inhibitor, 100X (Thermo Fisher: 1861280)

Spike in DNA for calibration (optional-- We use *Saccharomyces cerevisiae* micrococcal nuclease treated chromatin, originally provided by Henikoff lab)

RNAse A (Thermo Scientific: EN0531)

Glycogen (Thermo Scientific: R0561)

Bovine Serum Albumin (Sigma: A3059)

PCR purification kit (Qiagen, 28104)

DNA clean and concentrator (Zymo Research: D4033)

Qubit 1X dsDNA HS Assay kit (Thermo Fisher Scientific: Q33230)

**Equipment:**

Magnetic stand conFigureured for 1.5 ml tubes (Thermo Fisher Scientific, 12321D)

Thermomixer (Eppendorf, 5384000020)

Qubit 4 fluorometer (Thermo Fisher Scientific, Q33226)

Tube rotator (Grant-Bio, PTR-35)

Tube nutator (VWR, 82007-202)

**Solutions:**

**Digitonin (5%)** Dissolve 50 mg digitonin in 1 mL DMSO. Stable at room temperature for up to 1 week, or freeze at -20 °C.

**Binding buffer**

200 µL 1M HEPES pH 7.5

100 µL 1M KCl

10 µL 1M $CaCl_2$

10 µL 1M $MnCl_2$

Bring the final volume to 10 mL with ddH2O. Store the buffer at 4 °C for 6 months.

**Wash buffer**

1 mL 1 M HEPES pH 7.5

1.5 mL 5 M NaCl

12.5 µL 2 M spermidine

500 µL100X halt protease inhibitor

Bring the final volume to 50 mL with $dH_2O$. Store the buffer at 4 °C for up to 1 week.

**Dig-wash buffer**

400 µL 5% digitonin + 40 mL Wash buffer. Store the buffer at 4°C for up to 2 days.

**Antibody buffer**

8 µL 0.5 M EDTA

40.2 µL 5% BSA

2 mL Dig-wash buffer

Keep the buffer on ice.

**2X Stop buffer**

200 µL 4M NaCl

160 µL 0.5M EDTA

80 µL 0.2M EGTA

20 µL10mg/mL RNAse

32 µL 5 mg/mL Glycogen

16 µL 5% digitonin

2 µL 10 pg/µL yeast spike in DNA (optional)

Up to 4ml ddH20. Store the buffer at 4 °C for up to 1 week.

**Con-A bead activation:**

1. Gently resuspend the Concanavalin-A (Con-A) coated magnetic beads and withdraw a calculated total volume such that there will be 10 µL for each intended reaction. Place the total volume in a single 1.5mL eppi tube.

2. Wash beads by adding 1- 1.5 mL binding buffer to the eppi tube and gently pipetting up and down several times.

3. Place the tube on a magnetic stand and wait 1-2 minutes. When liquid appears clear, remove and discard.

4. After removing the tube from the magnetic stand, resuspend beads in 1.5 mL binding buffer and mix by gentle pipetting.

5. Return the tube to the magnetic stand, wait 1-2 minutes and when liquid appears clear, remove and discard.

6. After removing the tube from the magnetic stand, resuspend the beads in a volume of binding buffer equal to the initial volume of the bead suspension (step 1).

**Binding cells to activated beads:**

**NOTE:** We perform these steps at room temperature with room temperature buffers to minimize cell aggregates that cause bead clumping. We have found performing these steps with cold buffers results in cell aggregation that significantly reduces protocol efficiency.

7. Wash embryos with fish water in a petri dish, manually remove chorions with forceps, and transfer embryos to an eppi tube with wide-bore tip. Embryo number should be adjusted to yield a total of 50-100K cells per reaction (For 6hpf ~25 embryos-assuming 4,000 cells per embryo). At this stage, embryos for all intended reactions are pooled in a single 1.5 ml eppi tube. Remember to include reactions for IgG negative controls.

**NOTE:** Using a sharp forceps is critical for quick dechorionation. We use Finescience (11254-20) and can dechorionate ~100 embryo/10 min. Since stage match between different experimental groups is critical for an accurate interpretation, we recommend starting dechorionation early (4-5 hpf) and keeping dechorionated embryos at incubator until they reach the appropriate stage.

8. Remove as much fish water as possible from the embryos and gently wash whole embryos with 200 μL 1X PBS.

9. Remove the PBS, add 200 μL wash buffer, and gently pipette up and down with p200 to dissociate the embryos into a single cell suspension.  (It is critical that cells are in a single cell suspension).

10. Spin cells down at 600g for 3 min, remove supernatant from cell pellet.

11. Resuspend cell pellet in at least 1 ml of wash buffer at room temperature, centrifuge 3 min 600xg at room temperature and remove liquid.

12. Resuspend cell pellet in 1ml wash buffer.

**NOTE:** For ~500K cells (5 reactions), we add 1 mL wash buffer. Wash buffer can be adjusted depending on the total number of intended reactions.

13. While vortexing gently (1100 rpm), add the Con-A bead slurry dropwise. Place on end-over-end rotator at room temperature for 10 min.

14. After a quick spin to remove liquid from cap (<100xg), place the tube on magnet stand for 1-2 min. Remove liquid when it becomes clear.

**Antibody binding:**

15. Add 50 µL antibody buffer per reaction (ie: 250 µL for 5 reactions) to the tube containing the cells bound to Con-A beads. Very gently pipette up and down to fully resuspend beads.

16. For each reaction, aliquot 50 µL bead suspension to an individual 1.5ml eppi tube and add primary antibody (typically a 1:50-1:100 dilution, or 0.5-1 ul). Include negative control reactions (e.g. rabbit IgG).

17. Place the tubes on a nutator at 4°C and nutate overnight to several days (longer incubation times increase the reproducibility of the protocol). Alternatively, nutate for 2 hours at room temp. Be careful to make sure beads remain immersed in liquid and do not dry out.

18. Place each tube on the magnetic stand for 1-2 minutes and remove liquid when it appears clear.

**Binding of pAG-MNase:**

19. Add 0.8-1mL Dig-wash buffer to each tube. Gently pipette up and down to resuspend the beads.

20. Place the tube on the magnetic stand for 1-2 minutes and remove the liquid when it appears clear.

21. Repeat steps 19 & 20 for a total of two washes.

22. Add 40 µL Dig-wash buffer to the beads, pipetting along the side of the tube. Gently pipet to dislodge and resuspend beads.

23. Add 2 µL pAG-MNase (Epicypher) to beads resuspended in Dig wash buffer.

-**NOTE:** pAG-MNase plasmid is also available at Addgene (cat. no. 123461) and can be expressed and purified as previously described (Meers 2019).

24. Nutate the tubes on the nutator at room temperature for 10 min.

25. Place the tubes on a magnetic stand, wait 1-2 minutes and remove the liquid from each tube when it appears clear

26. Add 1 mL Dig-wash buffer to each tube, mix by gentle pipetting.

27. Return the tubes to the magnetic stand and remove Dig-wash buffer.

28. Repeat Dig-wash Steps 26 & 27 for a total of two washes.

**Targeted chromatin digestion:**

**NOTE:** It is critical to keep tubes at 0-4°C during the next steps to minimize background chromatin cleavage by the pAG-MNase.

29. For each tube, resuspend the beads in 100 µL of Dig-wash buffer. Gently pipette to dislodge and resuspend the beads.

30. Chill the tubes and 100 mM $CaCl_2$ on ice for ~30min.

31. Add 2 µL 100 mM $CaCl_2$ to each tube and mix with gentle vortexing (or pipetting). Immediately put the tubes on ice.

32. Nutate the tubes on a tube nutater for 2 hours at 4°C.

**Chromatin release:**

33. Add 100 µL of 2X STOP buffer to each tube and mix by gentle vortexing.

34. Incubate the tubes at 37°C for 10min on a ThermoMixer at 500 rpm to release fragments into solution.

35. Place the tubes on the magnet stand 1-2 minutes, waiting for liquid to appear clear. Cleanly transfer the supernatant containing digested chromatin to a fresh 1.5ml tube.

36. Clean and concentrate the DNA using a column or phenol-chloroform-isoamyl alcohol. For histone modifications, Qiagen PCR clean up columns provide good results, but columns with smaller size cutoffs such as Zymo Research DNA clean and concentrator may be required when examining transcription factors with smaller footprints.

37 OPTIONAL: Measure DNA concentration using Qubit assay.
This step is optional as getting low signals (~0.1 ng/µL) from the qubit assay does not necessarily mean that the CUT&RUN did not work, just that you recovered very little DNA. More often, we use positive bioanalyzer traces after the library preparation as a measure of quality control.

38. Keep the DNA samples at -20 °C until you are ready to prepare libraries.

#### B. CUT&RUN Library preparation

**NOTES:**

1. Using less than 30 ng DNA is recommended to prevent saturation of enzyme activities. CUT&RUN elute can be contaminated with large DNA fragments that result in high DNA concentration. Using bioanalyzer to quantify the relative amount of short DNA fraction can help with efficient library preparation.

2. Care should be taken during library preparation when dealing with CUT&RUN fragments below ~150 bp to avoid loss during purification steps. Additional optimization or alternative library preparation protocols may be required for very small fragments.

**Key Reagents:**

Sera-Mag SpeedBead Carboxylate-Modified Magnetic Particles (Hydrophobic), 15 mL Cat# 65152105050250

50 bp DNA ladder (cat#10416014)

Illumina iTruSeq Adaptor

i7 and i5 primers

NEB Ultra II End Repair Module (cat # E7546S)

NEBNext Ultra II Q5 Hot Start HiFi  PCR Master Mix (cat # M0543S)

T4 DNA ligase (M0202S)

**Solutions:**

**1X TE (50 ml)**

5 ml 1M Tris.Cl (pH 8)

100 µL 0.5M EDTA

**PEG solution (All will be used for 100 µL Sera-Mag beads):**

0.9 g PEG-800

1 ml 5M NaCl

2.5 ml TE (pH 8)

Vortex for 3-4 min until PEG goes into the solution.

**Preparing beads Sera-Mag Speed Beads**

1.Mix Sera-Mag speed beads and transfer 100 µL to a 1.5 ml eppi tube.

2. Place tube on the DynaMag-2 and let beads sit for 6 minutes on the magnet. Remove the clear supernatant (Use p200 pipette and do it slowly).

3. Add 1 ml 1X TE buffer pH 8. Remove the tube from the magnet and mix by pipetting. Place the tube on the magnet and let beads sit until the supernatant is clear. Discard the supernatant. Resuspend the beads in 100 µL 1X TE.

4. Add clean Sera-Mag beads to the PEG solution. To get the remaining beads from the tube, add 100 µL of TE buffer and transfer it into the bead/PEG solution. Bring volume up to 5 ml by adding TE buffer (~0.5-0.7 ml) and keep at 4°C.

**Testing beads**

**NOTE: Since beads ratio is critical for a correct size selection of DNA, test the beads monthly before use.**

1. Prepare fresh 3 ml 80% ethanol.

2. Mix 2 ul of generuler 50 bp DNA ladder with 18 ul of nuclease free water into each well of 8-strip tubes.

X volume of Beads / 20 ul of DNA ladder solution

| | | |
|---|---|---|
| Well #1, | control | (No beads) |
| Well #2, | 0.6 | (add 12 µl of beads) |
| Well #3, | 0.8 | (add 16 µl of beads) |
| Well #4, | 1 | ( add 20 µl of beads) |
| Well #5, | 1.2 | (add 24 µl of beads) |
| Well #6 | 1.4 | (add 28 µl of beads) |
| Well #7 | 1.6 | (add 32 µl of beads) |
| Well #8 | 2 | (add 40 µl of beads) |

3. Mix the beads and DNA ladder by pipetting. Incubate at room temperature for 10 min to allow the DNA binding to the beads.

4. Place the 8-strip tubes on a magnet and allow the beads to form a pellet. Remove clear supernatant except the control.

5. Perform ethanol washes on wells #2-8 (do not touch to the well #1). Add 200 µL 80% ethanol ,

and incubate for 30 seconds at room temperature while the beads are on magnet. Then carefully remove and discard the supernatant.

6. Repeat the previous step for 5 more times with a total of 6 ethanol washes.

7. Let beads dry at room temperature for ~10 min. (Be careful not to over dry the beads. They should be dried for no more than 10 min.)

8. Resuspend dry beads in 18 μl nuclease free water and keep them at RT for 10 min to elute the DNA.

9. Place tubes on the magnet. Transfer 15 μl of clear supernatant into an eppi tube and add 3 ul DNA loading dye.

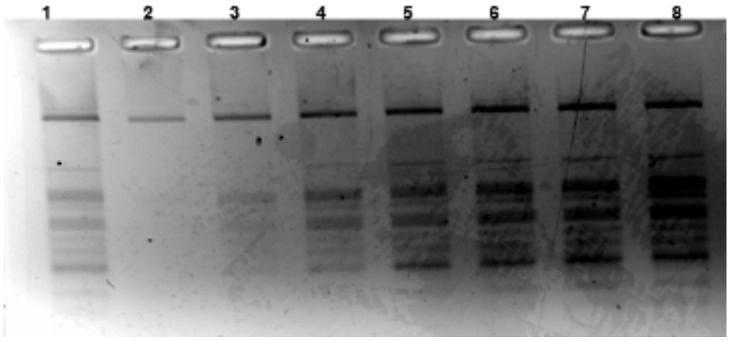


**Agarose gel image of size selected DNA ladders by different ratios of Sera-Mag Speed beads.** Well #2 (0.6X) picks up >500bp, Well #4 (1X), picks up >250bp and Well #6 (1.4X) picks up >150bp.

**<u>End repair</u>**

Perform End repair using NEB UltraII end repair module:

1. Mix:

   25.5 µl DNA

   3 µl 10X End repair reaction buffer

   1.5 µl End Prep Enzyme mix

2: Incubate:

   30 min at 20 ºC

   30 min at 65 ºC

   Hold at 4 ºC

**Adaptor Ligation**

1. Add the following directly to the end repair mix:

   4 ul 10X ligase buffer with dATP

   2 ul of T4 DNA ligase

   2 ul of iTruseq adaptor (1.5 uM stock)

   2 ul of ddH2O

   Total volume is 40 µl.

2. Mix by tapping. Do not vortex. Incubate overnight at 16 ºC

**Removal of self ligated adaptors**

1. Add 56 µl (1.4x, for histones) or 70 µl (1.75x, for Transcription factors) beads to the adaptor ligation reaction.

2. Incubate samples at room temperature for 5 min.

3. Place on the magnet for 5 min to clear the supernatant. Discard supernatant.

4. Add 200 μl freshly prepared 80% ethanol while beads were on the magnet, incubate for 30 sec and remove ethanol.

5. Repeat the ethanol wash one more time.

6. Air dry beads for 5 min on the magnet while the lids are open.

7. Remove tubes from the magnet and elute DNA in 22 μl of 10 mM Tris.Cl (pH 7.5-8).

8. Pipette the solution up and down. Incubate beads at room temperature for 2 min.

9. Place the tubes on the magnet and remove 20 μl of supernatant to a new PCR tube.

**PCR amplification**

using NEB ultra II Q5 hotstart polymerase (cat# MO543S)

1. Mix:

    20 μl Adaptor ligated DNA fragments.

    5 μl i7/i5 take 2.5 ul from individual stocks

    25 μl 2X Q5 Hot start polymerase master mix.

2. Amplify:

    Denature at 98 ºC 30 sec

    12 cycles of

98 ºC 10 sec

55 ºC 30 sec

72 ºC 60 sec

72 ºC 3 min (final extension)

Hold 10 ºC

OPTIONAL: Analyze the library: Quantify the library on qubit and run 10-20 ng on a gel to confirm correct size distribution.

**<u>Clean up of PCR amplification</u>**

**For Histones and other proteins with footprints >125 bp:**

1. Add 50 ul of beads (1x)

2. Incubate 5 min at room temperature.

3. Place on the magnet for 5 min to clear the supernatant.

4. Add 200 ul freshly prepared 80% ethanol into the tubes on the magnet, incubate 30 sec and remove ethanol.

5. Repeat the previous step.

6. Air dry beads for 5 min on the magnet with an open lid.

7. Remove tubes from the magnet and elute DNA in 15 ul of 10 mM Tris buffer.

8. Mix solution up and down. Incubate beads for 2 min at room temperature.

9. Spin quickly and place tube on the magnet for 5 min. Move 13 ul of supernatant to a new tube.

**Alternative method to improve recovery of small fragments (for use with Transcription factors)**

1. Add 40 ul beads (0.8x).

2. Incubate 5 min at room temperature.

3. Place the tube on the magnet to separate the beads from the supernatant.

4. After 5 min, carefully transfer the supernatant containing the DNA to a new tube. Discard the beads that contain the unwanted large fragments.

<u>Second round of size selection:</u>

5. Add 30 ul beads (1.4x) to the supernatant and pipette at least 10 times. Then incubate samples at room temperature for at least 5 min.

6. Place the tube on an appropriate magnetic stand for 5 min to separate the beads from the supernatant.

7. After the solution is clear, carefully remove and discard the supernatant. Be careful not to disturb the beads that contain DNA targets.

8. Add 200 ul of 80% freshly prepared ethanol to the tube on the magnet. Incubate at room temperature for 30 sec, then carefully remove and discard the supernatant.

9. Repeat the previous step one more time for a total of two washes. Be sure to remove all visible liquid after the second wash. If necessary, briefly spin the tube, place back on the magnet and remove traces of ethanol.

10. Air dry the beads while the tubes on the magnet stand with open lid.

11. Remove the tube from the magnetic stand. Elute the DNA target from the beads by adding 15 ul of 0.1x TE.

12. Mix well by pipetting up and down 10 times. Incubate for at least 2 min at room temperature.

13. Place the tubes on the magnetic stand. After 5 min (or when the solution is clear), transfer 13 ul to a new PCR tube and store at -20 ºC.

#### II. Protocol for CUT&Tag of zebrafish embryos

**Note:** This experimental protocol for performing CUT&Tag from nuclei using zebrafish embryos was adapted from Kaya-Okur, *et. al.*, Nature protocols, 2020.

**Buffers**:

**Bead Activation Buffer (211 uL per sample)**

200 µL 1M HEPES-KOH pH 7.9

100 µL 1M KCl

10 µL 1M $CaCl_2$

10 µL 1M $MnCl_2$

9.68 mL dd$H_2O$

**Nuclear Extraction Buffer (200uL per sample)**

1 mL 1M HEPES-KOH pH 7.9

500 µL 1 M KCl

12.5 µL 2 M spermidine

500 µL 10% (vol/vol) Triton-X100

10 mL glycerol

38 mL dd$H_2O$

1 Roche Complete Protease Inhibitor EDTA-Free tablet

**Wash150 Buffer**

1 mL 1 M HEPES pH 7.5

1.5 mL 5 M NaCl

12.5 µL 2 M spermidine

47.5 mL with $ddH_2O$

1 Roche Complete Protease Inhibitor EDTA-Free tablet

**Digitonin (5%)**

Dissolve 50 mg digitonin in 1 mL DMSO. Aliquot and store at -20 °C.

**Digitonin150 Buffer (450 uL per sample)**

499 uL Wash150 Buffer

1 uL 5% Digitonin

**Antibody150 Buffer (50 uL per sample)**

498 uL Digitonin150 Buffer

2 uL 0.5 M EDTA

**Wash300 Buffer**

1 mL 1 M HEPES pH 7.5

3 mL 5 M NaCl

12.5 μL 2 M spermidine

46 mL with $ddH_2O$

1 Roche Complete Protease Inhibitor EDTA-Free tablet

Digitonin300 Buffer (450 uL per sample)

499 uL Wash300 Buffer

1 uL 5% Digitonin

**Tagmentation Buffer (50 uL per sample)**

99 uL Wash300 Buffer

1 uL 1 M $MgCl_2$

**TAPS Buffer (50 uL per sample)**

5 mL 10 mM TAPS, pH 8.5

2 uL 0.5 M EDTA

**SDS Release Buffer (5 uL per sample)**

99 uL 10 mM TAPS, pH 8.5

1 uL 10% SDS

**SDS Quench Buffer (15 uL per sample)**

67 uL Triton-X 100 in 10mL $ddH_2O$

**<u>ConA bead activation</u>:**

1. Gently resuspend the Concanavalin A (ConA) coated magnetic beads and withdraw a calculated total volume (11 uL/sample) to ensure that there will be 10 μL for each intended reaction. Place the total volume in a single 1.5 mL eppi tube for batch processing.

2. Place the tube on a 1.5 mL magnetic stand and wait 1-2 minutes until slurry clears and pipet to remove supernatant.

3. After removing the tube from the magnetic stand, resuspend beads with 100 uL/sample cold Bead Activation Buffer, and mix by gentle pipetting. Place the tube on a magnetic stand until slurry clears and pipet to remove supernatant.

4. Repeat previous step for total of two washes.

5. Resuspend beads with 11 uL/sample cold Bead Activation Buffer.

6. Aliquot 10 uL/sample of activated ConA beads into 8-strip tube. Keep beads on ice until nuclei are ready.

**Nuclei preparation and binding nuclei to activated beads:**

7. Wash embryos with fish water in a petri dish, manually remove chorions with sharp forceps, and transfer embryos to an eppi tube with wide-bore tip. Embryo number should be adjusted to yield a total of 50-100K cells per reaction.

8. Remove as much fish water as possible from the embryos and gently wash whole embryos with 200 μL 1X PBS.

9. For 6 hpf embryos, remove the PBS, add 200 μL 1X PBS, and gently pipet up and down with p200 to dissociate the embryos into a single cell suspension. For 24hpf embryos, remove the PBS, add 200 uL 1X PBS, and pipet up and down with p200 to disrupt the yolk, spin down at 600g for 3 min, remove supernatant. Add protease solution containing 0.25% trypsin and 2 mg/mL Collagenase P in 1X PBS, and incubate at 28.5°C for 15-25 min, pipetting up and down every 5 min to aid cell dissociation into a single cell suspension.

10. Spin cells down at 600g for 3 min at room temperature, remove supernatant from cell pellet.

11. Resuspend cells in 100 uL/sample cold Nuclear Extraction Buffer, and pipet gently to mix. Incubate for 10 min on ice.

12. Spin nuclei down at 600g for 5 min at room temperature, remove supernatant from nuclei pellet.

13. Resuspend nuclei in 100 uL/sample cold Nuclear Extraction Buffer.

14. Aliquot 100 uL nuclei to each 8-strip tube containing 10 uL activated beads. Gently vortex to mix.

15. Incubate nuclei and beads on nutator for 10 min at room temperature.

**Primary and secondary antibodies binding:**

16. Place the tube on magnet stand for 1-2 min until slurry clears, and pipet to remove supernatant.

17. Add 50 uL cold Antibody150 Buffer to each sample, remove from magnet, and gently pipet up and down to thoroughly resuspend beads.

18. Add primary antibody to each tube (typically using 1:50-1:100 dilution), and gently vortex. Ideally to include a sample of IgG antibody (or without primary antibody of interest) as negative control, and a sample of H3K27me3 as positive control.

19. Incubate 8-strip tube on nutator at room temperature for at least 1 h or at 4°C overnight. Beads should not clump throughout the procedure.

20. Place the tube on a magnet until slurry clears and pipet to remove supernatant.

21. Remove from magnet, add 50 µL cold Digitonin150 Buffer to each 8-strip tube, and thoroughly pipet to resuspend.

22. Add secondary antibody (using 1:100 dilution, or 0.5 uL) to the tube, and gently vortex.

23. Incubate 8-strip tube on a nutator for 30 min at room temperature.

24. Place the tube on a magnet until the slurry clears and pipet to remove supernatant.

25. While beads are on magnet, add 200 µL cold Digitonin150 Buffer directly onto beads of each sample, and then pipet to remove supernatant.

26. Repeat the previous step for a total of two washes.

**Binding of pAG-Tn5:**

27. Add 2.5 uL/sample pAG-Tn5 (20X stock from EpiCypher) or home-made pAG-Tn5 to 50 uL/sample cold Digitonin300 Buffer, gently vortex to mix or pipet with p200 to mix to ensure thorough resuspension.

28. Incubate samples on nutator for 1 hr at room temperature, return 8-strip tube to magnet, and pipet to remove supernatant.

29. Add 200 µL cold Digitonin300 Buffer directly to each sample, thoroughly resuspend by pipetting, return to magnet, and then pipet to remove supernatant.

30. Repeat the previous step for a total of two washes.

**Targeted chromatin tagmentation:**

31. Remove from magnet, add 50 µL cold Tagmentation Buffer to each sample, and thoroughly pipet to resuspend.

32. Incubate 8-strip tube for 1 hr at 37°C in thermocycler for tagmentation.

33. Place the tube on a magnet until slurry clears and pipet to remove supernatant.

34. Remove from magnet and resuspend beads in 50 µL room temperature TAPS Buffer by pipetting, return to magnet, and pipet to remove supernatant.

35. Remove from magnet, add 5 µL room temperature SDS Release Buffer (containing 0.1% SDS) to each sample, and vortex on max speed for 7 seconds. Quick spin to collect.

36. Incubate 8-strip tube for 1 hr at 58°C in thermocycler.

37. Add 15 µL room temperature SDS Quench Buffer (containing 0.67% Triton-X) to each sample, and vortex on max speed.

**Library preparation and cleanup**:

38. For each sample, add 2 µL each of barcoded i5 and barcoded i7 primers (10 µM stocks).

39. Add 25 uL non-hot start CUTANA High Fidelity 2x PCR Master Mix (EpiCypher) or NEBNext High-Fidelity 2X PCR Master Mix (NEB #M0541S) to each sample and mix.

40. Amplify based on the following PCR setting:

58 ºC 5 min

72 ºC 5 min

98 ºC 45 sec

14-21 cycles of

98 ºC 15 sec

60 ºC 10 sec

72 ºC 1 min

Hold 4 ºC

41. DNA cleanup using 1.3x AMPure beads to sample volume according to manufacturer's recommendations.

42. Elute DNA in 15 uL 0.1x TE buffer, and transfer to a new PCR tube and store the library at -20 ºC.
